# 3D spatially-resolved geometrical and functional models of human liver tissue reveal new aspects of NAFLD progression

**DOI:** 10.1101/572073

**Authors:** Fabián Segovia-Miranda, Hernán Morales-Navarrete, Michael Kücken, Vincent Moser, Sarah Seifert, Urska Repnik, Fabian Rost, Alexander Hendriks, Sebastian Hinz, Christoph Röcken, Dieter Lüthjohann, Yannis Kalaidzidis, Clemens Schafmayer, Lutz Brusch, Jochen Hampe, Marino Zerial

## Abstract

Early disease diagnosis is key for the effective treatment of diseases. It relies on the identification of biomarkers and morphological inspection of organs and tissues. Histopathological analysis of human biopsies is the gold standard to diagnose tissue alterations. However, this approach has low resolution and overlooks 3D structural changes that are consequence of functional alterations. Here, we applied multiphoton imaging, 3D digital reconstructions and computational simulations to generate spatially-resolved geometrical and functional models of human liver tissue at different stages of non-alcoholic fatty liver disease (NAFLD). We identified a set of new morphometric cellular parameters correlated with disease progression. Moreover, we found profound topological defects in the 3D bile canaliculi (BC) network. Personalized biliary fluid dynamic simulations predicted an increased pericentral biliary pressure and zonated cholestasis, consistent with elevated cholestatic biomarkers in patients’ sera. Our spatially-resolved models of human liver tissue can contribute to high-definition medicine by identifying quantitative multi-parametric cellular and tissue signatures to define disease progression and provide new insights into NAFLD pathophysiology.

## Introduction

High definition medicine is emerging as an integrated approach to profile and restore the health of an individual using a pipeline of multi-parametric analytical and therapeutic technologies^1^. High-definition medicine relies on large data sets, e.g. genomics, metabolomics, to characterize human health at the molecular level. It also relies on imaging, image analysis and computational modelling approaches to identify structural and functional abnormalities in organs and tissues associated with a disease state. Histology has been classically used to characterize tissue structure and remains the method of choice to describe and monitor a large variety of pathologies^2^. However, this technique has several disadvantages, e.g. it is subjective (depends on the pathologist’s skills), is often semi-quantitative and provides only two-dimensional (2D) information, i.e. does not account for the three-dimensional (3D) complexity of tissues^3^. In recent years, an increasing number of studies have highlighted the importance of considering 3D information for the histopathological examination of tissues^4-7^. This is particularly crucial for the analysis of 3D structures. The liver is a pertinent example of an organ with a complex 3D tissue organization^8^. It consists of functional units, the liver lobule^9,10^, containing two intertwined networks, the sinusoids for blood flow and the bile canaliculi (BC) for bile secretion and flux^9^. Sinusoids and BC run antiparallel along the central vein (CV)-portal vein (PV) axis. The hepatocytes are the major parenchymal cells and display a peculiar and unique type of cell polarity distinct from that of simple epithelia^11^. Whereas in epithelia all cells share the same orientation with their apical surface facing the lumen of the organ, hepatocytes are sandwiched between the sinusoidal endothelial cells and share the apical surface with multiple neighbouring hepatocytes to form a 3D BC network^12,13^. Such an architecture makes it difficult to grasp the 3D organization of cells and tissue from 2D histological sections. Recent advances in tissue staining, optical clearing and multi-photon microscopy allow imaging thick sections of tissues such that 3D information can be captured^14,15^. Computer software process the images to generate 3D digital reconstructions of tissues, i.e. geometrical models^8^, with single-cell resolution. These provide a detailed quantitative description of the different cells and structures forming the tissue. The geometrical information extracted from the 3D reconstruction can also be used to generate predictive models of tissue function e.g. biliary fluid dynamic^16^, thus gaining novel insights into liver tissue organization and function. Thereby, geometrical models can be used to quantitatively describe the tissue architecture and function, improving our understanding of liver biology and pathobiology.

Non-alcoholic fatty liver disease (NAFLD), defined as an accumulation of triglycerides and lipid droplets (LD) in the liver in absence of alcohol intake (infiltration in > 5% of hepatocytes), is rising to the most common chronic liver disease worldwide^17^. NAFLD includes a spectrum of liver diseases, ranging from simple steatosis to non-alcoholic steatohepatitis (NASH)^17^. Whereas steatosis is considered as a “non-progressive” status of the disease, NASH has the potential to progress to more severe stages, such as cirrhosis or hepatocellular carcinoma, leading eventually to liver failure and transplantation^3,17^. Thus, the understanding of the transition from steatosis to NASH as a disease-defining moment for NAFLD prognosis is key to a deeper understanding of disease pathophysiology. Liver biopsy and histological inspection of thin tissue slices (< 10 µm) constitute the current gold standard for the diagnosis of steatosis and NASH^2,17,18^. Unfortunately, due to the limitations in providing 3D information, alterations in 3D tissue structures such as BC^3,17^, have been overlooked. Therefore, new approaches are required to gain a more complete understanding of NAFLD establishment and its progression to NASH. In this study, we generated 3D spatially resolved geometrical and functional models of human liver tissue for different stages of NAFLD to contribute to a high definition medical diagnosis of disease establishment and progression.

## Results

### Tri-dimensional geometrical models of human liver tissue

To identify parameters that can quantitatively discriminate the transition from simple steatosis to early NASH (eNASH) we stained, imaged and digitally reconstructed human liver tissue in 3D. We focused on cell and nuclear morphology, LD, and tissue features such as BC and sinusoidal networks, and their spatial distribution along the CV-PV axis. For this we tested 26 antibodies combinatorially with dyes and antigen retrieval protocols (Supplementary Table 2). Our final pipeline (see Methods for details) includes the following steps. First, 100 µm liver sections were heated and antigen retrieved using citric acid buffer (CAAR). Second, we stained for BC (CD13), sinusoids (fibronectin), nuclei (DAPI), lipid droplets LD (BODIPY) and cell borders (LDLR), optically cleared with SeeDB^19^ and imaged at high resolution using multiphoton microscopy (0.3 µm x 0.3 µm x 0.3 µm per voxel) (Extended Data Fig. 1). Because this protocol did not provide a good cell border staining of the pericentral hepatocytes in STEA and eNASH, for cell-based measurements (see below Fig. 3), we used an alternative protocol without antigen retrieval enabling the staining of sinusoids (fibronectin), nuclei (DAPI), LD (BODIPY) and cell borders (phalloidin). We applied this pipeline to biopsies from twenty-two patients classified into four groups: normal control (NC, n = 6), healthy obese (HO, n = 4), steatosis (STEA, n = 5) and early NASH (eNASH, n = 7). The demographic, clinical and histological details of the samples are summarized in the Supplementary Table 1. All images cover one complete CV-PV axis within a liver lobule. Finally, we reconstructed the various stained components of the tissue using our open-source software Motion Tracking (http://motiontracking.mpi-cbg.de) as described^8^ (Fig. 1 and Supplementary Video 1-2). The generation of geometrical models constituted the basis for the quantitative and structural characterization of the different components forming the liver tissue in the NAFLD biopsies.

**Fig. 1.**
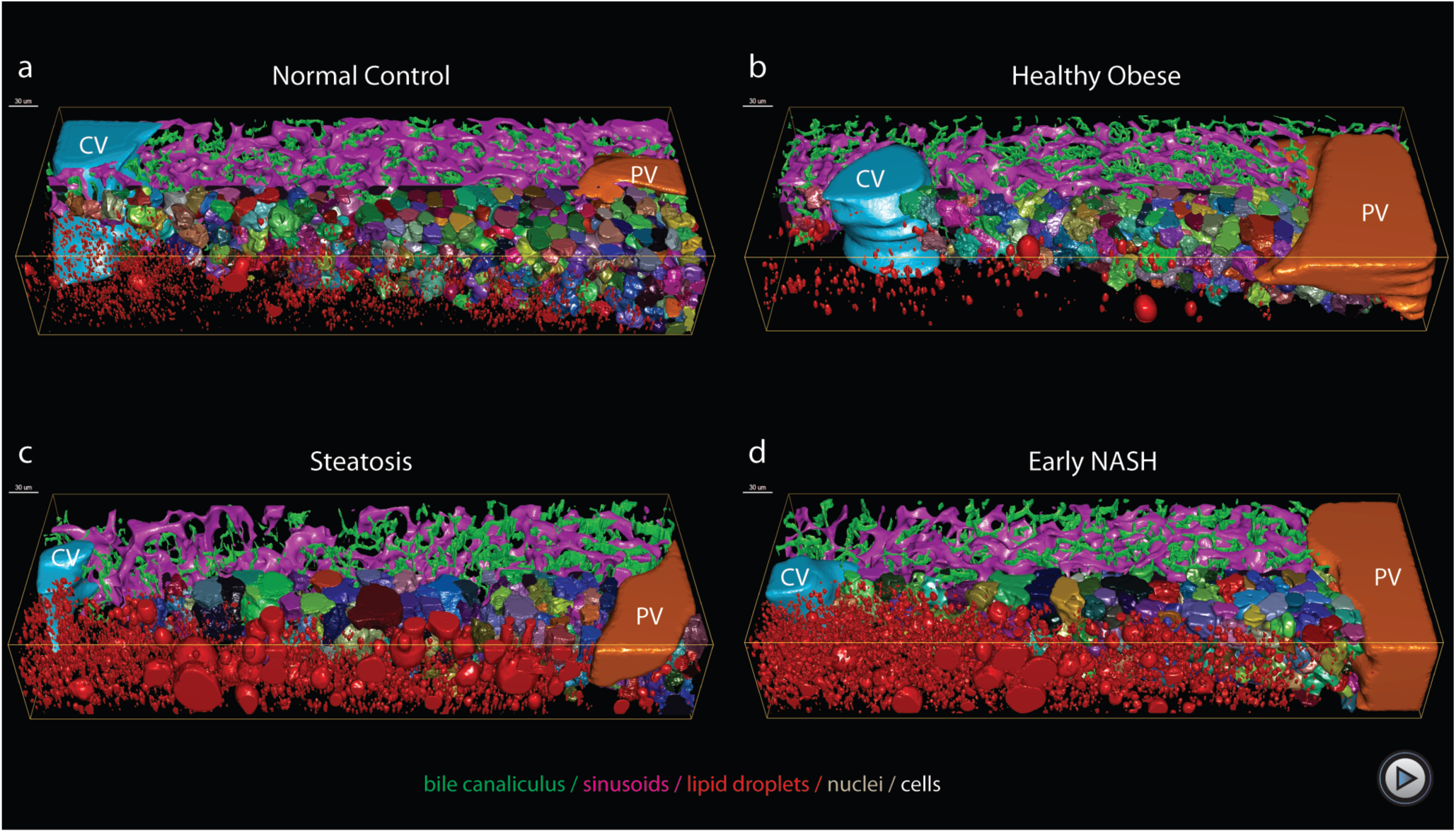
3D reconstruction and quantitative analysis of human liver morphology. Human liver sections obtained by biopsy (∼100 um thick) were stained for bile canaliculi (CD13), sinusoids (fibronectin), nucleus (DAPI), lipid droplets (BODIPY) and cell border (LDLR), optically cleared with SeeDB and imaged at high resolution using multiphoton microscopy (0.3 µm x 0.3 µm x 0.3 µm per voxel). For each sample, we reconstructed the central vein (light blue), portal vein (orange), bile canaliculus (green), sinusoids (magenta), lipid droplets (red), nuclei (random colours) and hepatocytes (random colours). **a**, Normal control. **b**, Healthy obese. **c**, Steatosis. **d**, Early NASH.

### Nuclear-based analysis of NAFLD progression

We first quantified properties of hepatocytes nuclei, such as cell nuclearity and ploidy, since hepatocytes are heterogenous in both mouse and human^8,20^. We quantified nuclear vacuolization/glycogenation given it is a common histological characteristic in NAFLD linked to insulin resistance^21,22^. Finally, we measured nuclear texture homogeneity, a feature associated with various pathological conditions, such as cancer, inflammation, cardiomyopathy, etc^23-26^, with methylation and acetylation status^27^ and, more recently, with transcriptional activity^28^. We neither observed differences in the proportion of mono/binuclear cells nor in the ploidy between the groups (Extended Data Fig. 2a and b). The average values of several nuclear features showed only modest variations Extended Data Fig. 2). However, many functional and morphological features of the liver change along the CV-PV axis, such as metabolic zonation^29,30^, ploidy^8^, cell volume^8^, BC^16^. Therefore, to account for potential morphological heterogeneities along the CV-PV axis, we computationally divided this axis into ten equidistant zones (Extended Data Fig. 2c) (Methods). We found major differences in nuclear elongation around the CV and vacuolization around the PV between the different groups (Extended Data Fig. 2d,e). Moreover, we identified zonated and progressive differences in nuclear homogeneity as disease progresses (Extended Data Fig. 2f-i). Therefore, our analysis reveals that, in spite of modest changes in the average values, the combined zonated values of nuclear vacuolization and texture homogeneity allow discriminating between all four patient groups.

### Morphometric parameters of LD correlate with disease progression

The finding that quantitative spatially-resolved analysis of nuclear parameters can reveal changes that are not evident in an average estimate prompted us to re-evaluate the morphometric characterization of LD. Even though LD are the most typical hallmark of NAFLD, a detailed quantitative description of their size and their spatial localization within the liver lobule has not been achieved yet. Contrary to traditional histology^3^, immunostaining of thick tissue sections preserved most of the LD (Fig 2a). In agreement with the histopathological description of NAFLD^3,17^, a major increase in LD was observed in STEA and eNASH, which were concentrated between the second and the fifth zones (Fig. 2b). However, the LD occupy a higher volume of the tissue in eNASH than STEA. It is known that the LD can present massive differences in size^31,32^. The LD in the human liver samples ranged from ∲1 µm^3^ to 20,000 µm^3^ (Fig. 2c). To inquire whether differences in LD size may correlate with the disease state, we performed a population analysis based on their volume distributions (Fig. 2c). We defined three sub-populations of LD, namely, small (< 14 µm^3^), medium (14 – 400 µm^3^) and large (> 400 µm^3^) ones (Fig. 2d-f). Strikingly, we found that the three sub-populations varied between disease conditions. Whereas small LD were evenly distributed along the CV-PV in all conditions, large LD were highly enriched towards the pericentral zone in STEA and eNASH, occupying up to 25% of the tissue volume (Fig. 2f). Most importantly however, medium LD were mostly present in eNASH and spanning the entire liver lobule (Fig. 2e), hence contributing to discriminate between STEA and eNASH in the periportal area (Fig. 2e).

**Fig. 2.**
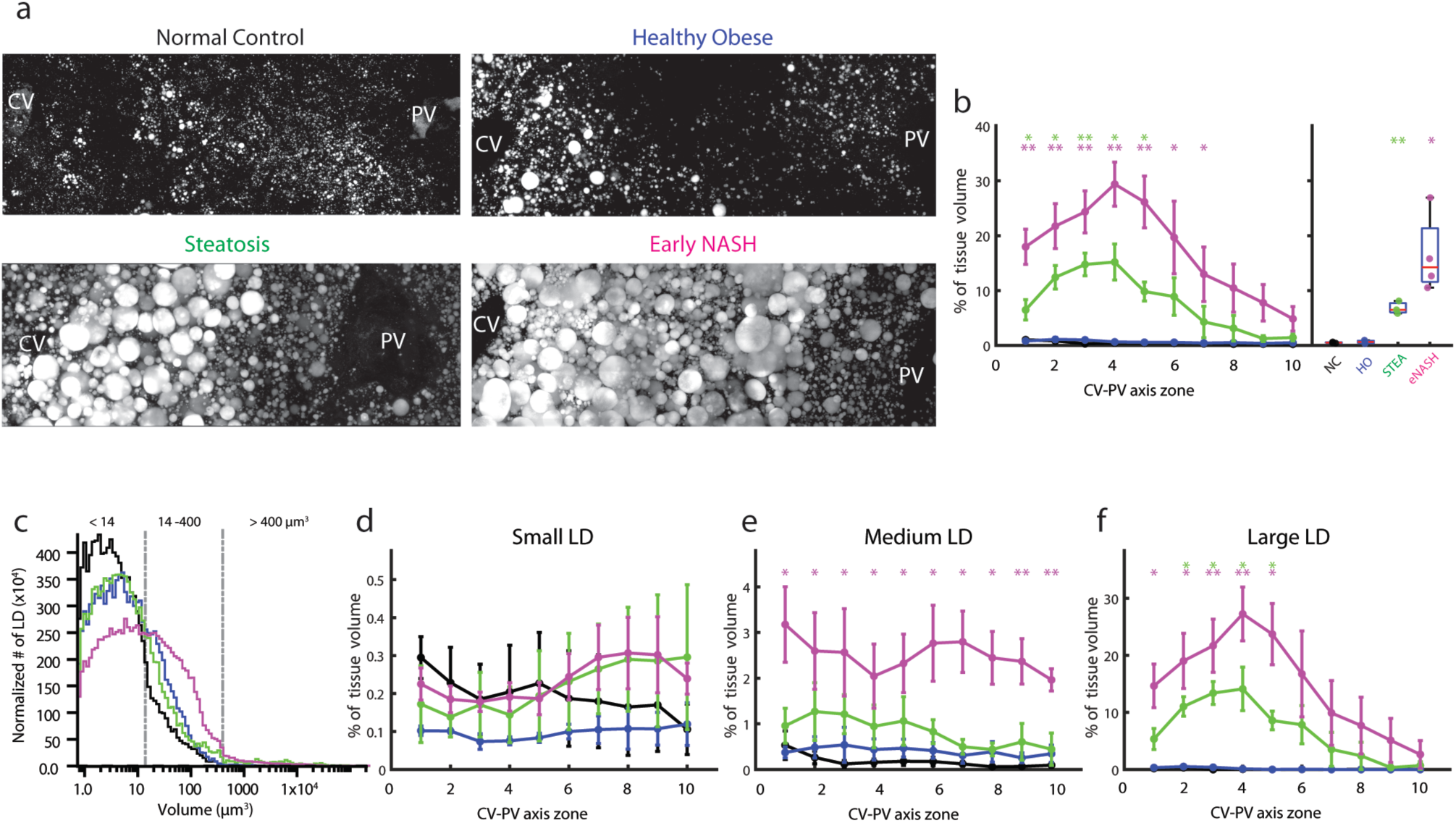
Quantitative characterization of LD along the CV-PV axis. **a**, Representative IF images of fixed human liver tissue sections stained with BODIPY. Shown is a maximum projection of a 60 µm z-stack covering an entire CV-PV axis. **b**, Quantification of the percentage of tissue volume occupied by the LD along the CV-PV axis and the overall values (i.e. over the whole CV-PV axis). **c**, LD volume distribution. LD populations were defined based on the LD volume distribution of the NC group (black line). Three populations of LD were defined based on their volume: small (< 14 µm^3^), medium (14 – 400 µm^3^) and large (> 400 µm^3^), corresponding to changes in the volume distribution (i.e. drops in the number of LD). Quantification of the percentage of tissue volume occupied by the LD along the CV-PV for (**d**) small, (**e**) medium and (**f**) large LD. NC = 3 samples, HO = 3 samples, STEA = 3 samples, eNASH = 3 samples. Spatially-resolved quantification represented by mean ± SEM per zone and overall quantifications by box-plots. *p-values < 0.05, **p-values < 0.01, ***p-values < 0.001.

The zonated increase in LD size during disease progression is such that some LD in the pericentral zone become even larger than a normal hepatocyte. This leads to global changes in tissue structure. To quantify such changes, we measured the spatial distribution of cell density, cell volume and percentage of cell volume occupied by LD. We found ∼50% reduction in the number of hepatocytes located between the CV and the middle zone in STEA and eNASH, when compared with NC and HO samples (Fig. 3a). This reduction was compensated by a massive increase in cell volume (Fig. 3b). Hepatocytes were two times larger than the average size (Fig. 3b), reaching values up to up to ∲100,000 µm^3^ for STEA and NASH (ten times bigger than a small hepatocyte) (Fig. 3c). A population analysis of the hepatocytes based on their volume revealed a characteristic distribution of different cells populations along the liver lobule (Fig. 3c-f). STEA and eNASH were characterized predominantly by small and large hepatocytes which are anti-correlated along CV-PV axis (Fig. 3d-f). Even though cell density and cell volume were practically indistinguishable between STEA and eNASH, we observed a remarkable phenotype regarding the fraction of cell volume occupied by LD (Fig. 3g-i). In eNASH, hepatocytes accumulated LD even in the periportal zone (Fig. 3h, 3i and Supplementary Video 3), suggesting that LD accumulation progressively extends to the PV as the disease progresses.

**Figure 3.**
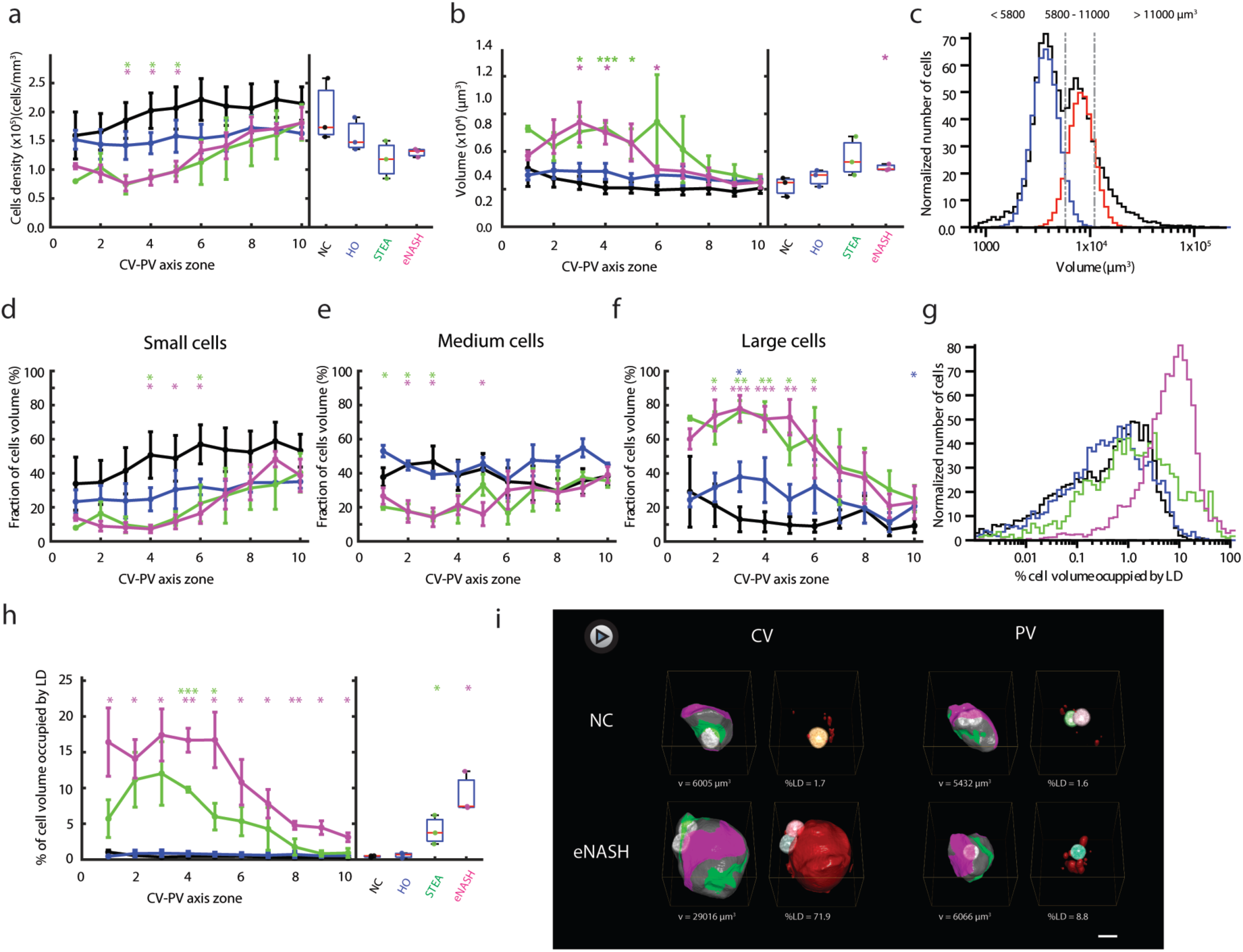
Cell based analysis of NAFLD. Quantification of the number of hepatocytes per tissue volume unit (**a**) and cell volume (**b**) both, along the liver lobule and the overall average. **c**, Cell volume distribution. For the population analysis, the hepatocytes form all the groups were pulled together and the populations were defined based on their volume distribution (**c**). By fitting the volume distribution with two normal distributions (**c**), the volume values defining three population’s boundaries were identified: small (< 5800 µm^3^) (**d**), medium (5800 – 11000 µm^3^) (**e**) and large (> 11000 µm^3^) (**f**). The fraction of cellular volume occupied by the different populations is shown in **d**, **e**, and **f**. Percentage of the cell volume occupied by lipid droplets: distribution (**g**) and statistics along the CV-PV axis and overall (**h**). NC = 3 samples, HO = 3 samples, STEA = 3 samples, eNASH = 3 samples. Spatially-resolved quantification represented by mean ± SEM per zone and overall quantifications by box-plots. *p-values < 0.05, **p-values < 0.01, ***p-values < 0.001. Representative cells reconstructed in 3D and selected from zone 3 and 8. Apical, basal and lateral surface are shown in green, magenta and grey respectively. Lipid droplets are shown in red. Scale bar, 10 µm.

Altogether, these data reveal profound quantitative morphological disparities in cell size and LD content along the CV-PV axis between NC, HO, STEA and eNASH. Specifically, the percentage of cell volume occupied by LD in the PV and CV zones can serve to discriminate STEA and eNASH.

### Alterations in apical protein trafficking

The massive presence of LD that occupy a large portion of the cytoplasm raises the question of whether trafficking of proteins to the apical plasma membrane of hepatocytes is affected. We analysed the localization of four apical proteins, aminopeptidase N (CD13), bile salt export pump (BSEP), multidrug resistant-associated protein (MRP2) and dipeptidylpeptidase 4 (DPPIV). CD13, BSEP and MRP2 were correctly localized to the apical membrane in all conditions (Extended Data Fig. 1, 3a-b). DPPIV was enriched on the apical membrane with a small fraction on the basal membrane in NC and HO (Extended Data Fig. 3c). Strikingly, DPPIV was redistributed to the lateral membrane in pericentral hepatocytes in STEA and eNASH, whereas it retained its normal localization on the periportal zone (Extended Data Fig. 3c). Considering that DPPIV follows the transcytotic route to the apical surface^33,34^ whereas BSEP and MRP2 do not^34-36^, the mislocalization of DPPIV suggests a possible disruption of some stage of endocytosis and transcytosis of this cargo molecule in the pericentral hepatocytes. This supports previous findings regarding the misregulation of membrane protein trafficking in NAFLD^37^ and prompted us to evaluate whether the integrity of the BC could be affected during the disease progression.

### Bile canaliculi network shows geometrical and topological-zonated defects in NAFLD

To determine whether 3D structures such as the BC and sinusoidal networks are affected, we carried out a geometrical and topological characterization of both networks. Even though we observed a slight reduction in the total length of the sinusoidal network in STEA and eNASH (Extended Data Fig. 4e), no major defects in sinusoidal microanatomy were detected (Extended Data Fig. 4a-d, f-g) (volume fraction, radius, branching and connectivity). Next, we analysed the BC network. Contrary to the very packed and homogeneous appearance in NC and HO, the BC in STEA and eNASH displayed clear morphological defects which were more pronounced in the pericentral zone (Fig. 4a, b). A more detailed analysis revealed a sustained increase in BC radius in NASH throughout the CV-PV axis (Fig. 4d). In addition, in both STEA and eNASH, we observed a strong reduction in the total length of the BC towards the pericentral zone (Fig. 4a and f). Other geometrical properties of the BC network, such as volume fraction and junction density were unaffected (Fig. 4c and e).

**Figure 4.**
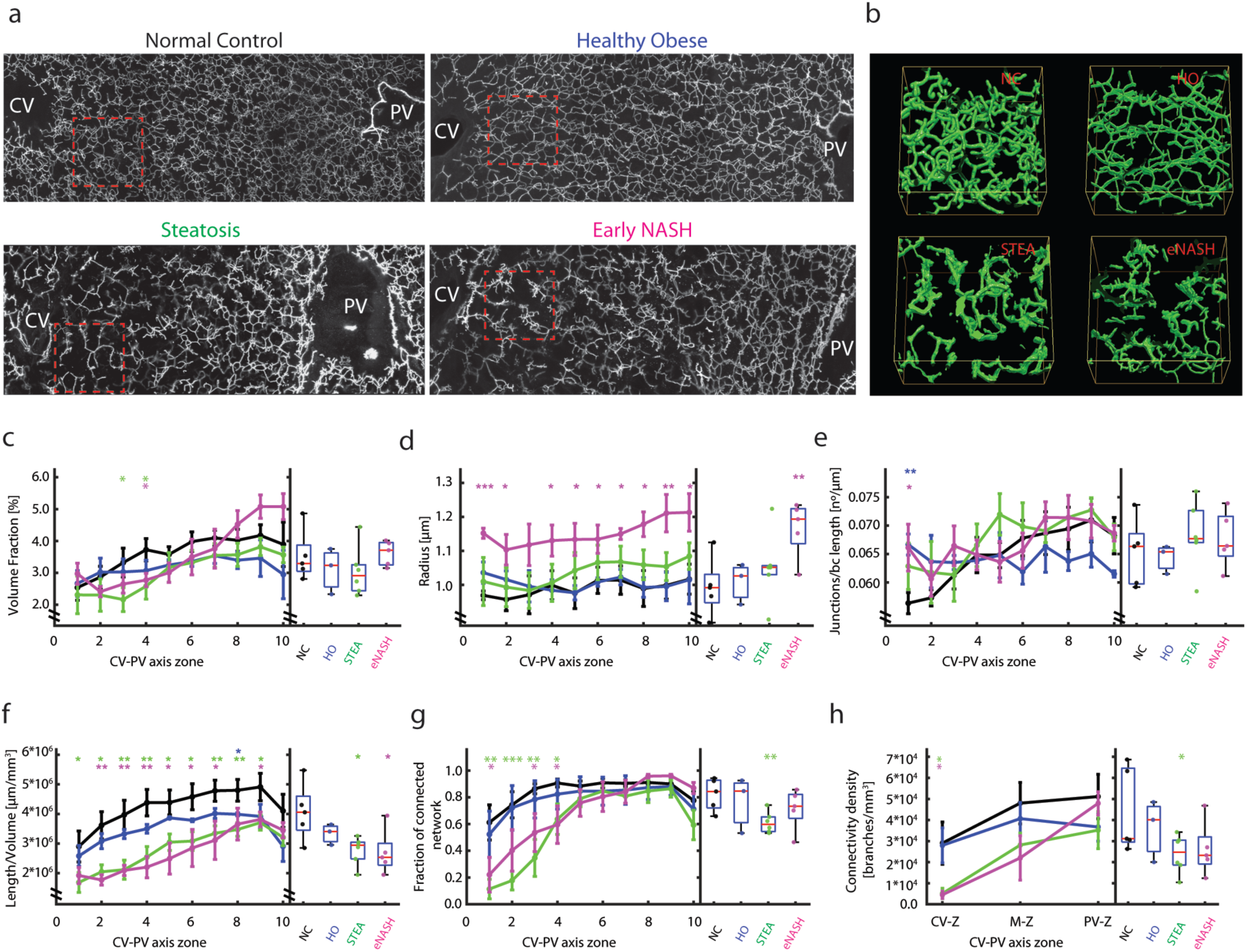
Structural and topological defects of bile canaliculi revealed by spatial 3D analysis. **a**, Representative IF images of fixed human liver tissue sections stained with CD13 after citric acid antigen retrieval. Shown is a maximum projection of a 60 µm z-stack covering an entire CV-PV axis. **b**, Inset showing 3D representation of the bile canaliculus highlighted in **a**. Quantification of the volume fraction of tissue occupied by bile canaliculi(**c**), radius (**d**), number of junctions (**e**), total length per volume (**f**), fraction of connected network (**g**) and connectivity density (**h**) of the BC network along the CV-PV axis and overall (See Methods for details). NC = 5 samples, HO = 3 samples, STEA = 6 samples, eNASH = 5 samples. Spatially-resolved quantification represented by mean ± SEM per zone and overall quantifications by box-plots. *p-values < 0.05, **p-values < 0.01, ***p-values < 0.001.

Finally, to investigate the topological properties of the BC network, we performed an analysis of network connectivity (see Methods). Surprisingly, we found a pronounced decay in the connectivity in STEA and eNASH towards the pericentral region (Fig. 4a, b, g and h, Supplementary Video 4). One possibility is that the alterations in BC may be the consequence of the spatial constrains arising from the presence of large cells in the tissue (Fig. 3f). To test this, we inspected the connectivity of the sinusoidal network. We found that no defect was observed for the sinusoidal network (Extended Data Fig. 4a, f and g), supporting the idea that the BC network is specifically affected and not an indirect consequence of spatial constraints. Thus, our data point at specific geometrical and topological alterations in the pericentral BC network in both STEA and eNASH.

### Personalised model of bile flow predicts increase in bile pressure in the pericentral zone

The observed alterations of BC network architecture are likely to have consequences for liver tissue function, particularly for bile flow. Clearly, information on bile velocity and pressure in disease conditions could be insightful. However, it is not yet possible to measure bile flow in the human liver at the level of BC as in animal models. We recently developed a computational model of bile fluid dynamics, validated its quantitative predictions in mouse models and demonstrated that bile velocity and bile pressure distributions along the liver lobule strongly depend on BC geometry^16^. However, this model^16^ is not suitable to handle the extreme inhomogeneity of BC density such as the ones encountered in tissue distorted by the presence of large LD (i.e. in STEA and eNASH). Therefore, we addressed this issue and further developed our model in a spatially-resolved fashion (Fig. 5a). Shortly, the refined model is based on conservation of mass for water and osmolytes and Darcy’s Law for laminar flow. The proportionality constant in Darcy’s Law was derived from the porous media theory. Boundary conditions were set to zero velocity at the outer surface of the central vein, and ambient pressure at the portal outlet. Since we obtained morphometric data for liver tissue from individual patients, we aimed at developing personalised models, i.e. parameterized by individual geometrical measurements (BC volume fraction *ε*_BC_, BC radius *r*_BC_., fraction of connected BC, canaliculi tortuosity τ, apical surface density *A*, intra-canalicular volume fraction occupied by microvilli 1-α) (Fig. 4c, 4d, 4g, and Extended Data Fig. 5) and previously reported values (viscosity, permeability and osmolyte secretion rate). No free parameters remained and, hence, no parameter fitting was needed (see Methods for details). Next, we applied this model to predict bile velocity, pressure and solute concentration distributions across the liver lobule for individual patients liver tissue 3D reconstructions from all the four groups.

**Figure 5.**
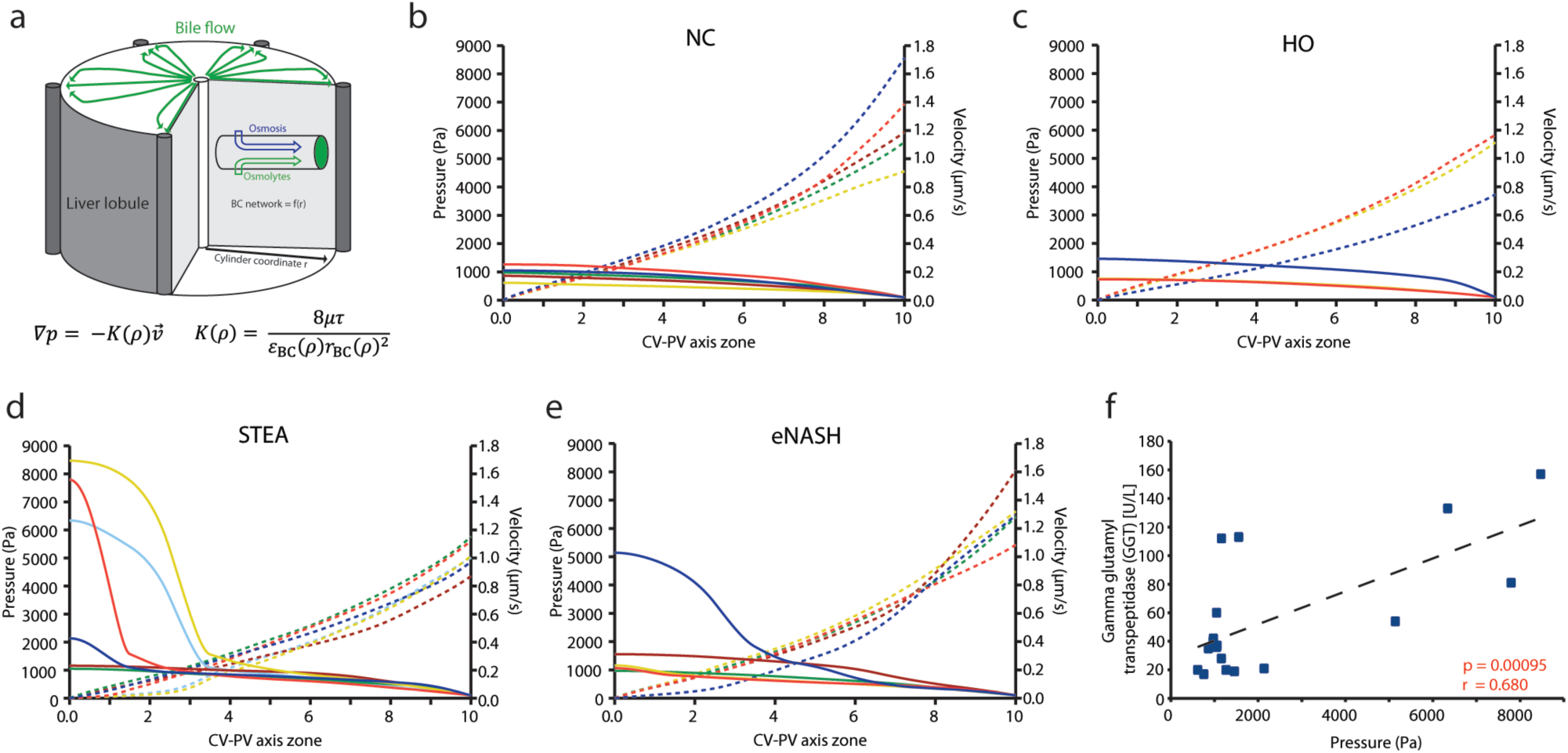
Individual-based model prediction of bile pressure *p* and flow velocity 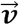 profiles based on measured bile canalicular geometries. a, **a**, Abstraction of liver lobule by cylinder symmetry with radial coordinate ρ. The mechanistic model considers secretion of osmolytes (green) and osmotic water influx (blue) in a porous medium with ρ-dependent properties (see supplemental model description). Darcy’s law is assumed with a proportionality constant K(ρ) depending on viscosity *μ*, tortuosity *τ*, bile canalicular volume fraction *ε*_BC_, bile canalicular radius *r*_BC_. All geometric parameters have been measured per patient. **b-e**, Model prediction for bile fluid pressure (solid line, left axis) and bile flow velocity (dashed line, right axis) profiles for individual patients (colour) in for disease groups. NC = 5 samples, HO = 3 samples, STEA = 6 samples, eNASH = 5 samples. **f**, Scatter plot of measured Gamma glutamyl transpeptidase (GGT) levels versus predicted pericentral (zone 0) bile fluid pressure from individual patients from all groups reveals a statistically significant positive correlation. One-sided t-test. P-values and Pearson correlation coefficient are indicated in the plots.

The model predicts bile velocities in the periportal area of about 1.2 ± 0.4 μm/sec for all patient groups (Fig. 5b-e). Very similar velocities have been reported in mouse^16^. However, the predicted pressure in the pericentral area differed significantly between the patient groups. In the NC and HO groups this pressure was predicted to be lower than 1500 Pa in all patients (963.2 ± 285.2 Pa, mean ± SD) (Fig. 5b-c). This is consistent with the reported maximum biliary secretion pressure of 1,000–1,500 Pa in the extrahepatic biliary system in rats^38^. In the STEA and eNASH groups, the model predicted an abrupt increase of bile pressure toward the pericentral zone (Fig. 5d-e). For six STEA and eNASH patients (55%), the pericentral pressure exceeded 1,500 Pa and for four patients (36%) it exceeded twice the maximum pressure predicted for NC (3000 Pa) (Fig. 5d-e). The compensatory effect in bile pressure observed in eNASH is mostly due to the dilation of BC (Fig. 4d). Therefore, our model predicts an increase in pericentral bile pressure in STEA and eNASH conditions ranging from relatively mild to quite severe, depending on the BC geometry of individual patients.

We next set out to validate the model predictions. As it is impossible to measure bile flow and pressure in the human liver, we considered possible consequences of changes in bile pressure. Increased bile pressure is a hallmark of cholestasis^39,40^. Therefore, as readout of increased bile pressure, we analysed the most commonly used cholestatic biomarkers in serum, including bilirubin, gamma glutamyl transpeptidase (GGT), alkaline phosphatase (ALP) and BAs. To increase the statistical power, we analysed additional sera samples for the different groups (NC = 25, HO = 25, STEA = 24 and eNASH = 26 samples). Whereas we found that both bilirubin and GGT were elevated in STEA and eNASH (Extended Data Fig. 6a-b), we did not detect significant changes in the levels of ALP, total BAs and primary BAs between the groups (Extended Data Fig. 6c, 7b,d). Strikingly, when we analysed the correlation between the predicted pericentral bile pressure and the biomarkers for individual patients from all groups, we found a strong correlation for the majority of the cholestatic biomarkers, with GGT having the strongest correlation (Pearson correlation coefficient ALP= 0.473, total BAs 0.505, primary BA 0.518 and GGT 0.680; (Extended Data Fig. 8a-e and Fig. 5f). In contrast, aspartate aminotransferase (AST) and alanine aminotransferase (ALT), biomarkers of hepatocellular liver damage which are not increased in cholestasis, showed no correlation (Extended Data Fig. 8f-g). The presence of pericentral cholestasis was also supported by the increase in the predicted pericentral concentration of BAs (proportional to the lumped concentration of all osmolytes. Extended Data Fig. 9), which in combination with high pressure and low flow velocity, are signs of a defective bile flow. Altogether, our model predicts a significant degree of zonated cholestasis as a new component of the NAFLD pathophysiology.

## Discussion

High definition medicine provides a novel approach to understand human health of individuals with unprecedented precision^1^. One of its pillars is the combination of image analysis and computational modelling to uncover tissue alterations at different structural and functional levels during disease progression. During the last years, there has been an enormous interest in getting a better understanding of NAFLD establishment and progression due to its growing impact on public health^41^. A lot of attention has been mostly drawn to the role of signalling pathways^42-44^, microbiome^45,46^, metabolism^47^, genetic risk factors^48^, BAs^44^, etc. However, a major challenge is to understand how the molecular alterations detected are expression of the organ dysfunction, manifested as morphological and functional alterations of cells and tissue architecture.

The classical histological analysis has provided insights into fundamental aspects of NAFLD. However, a quantitative description of the 3D tissue morphology is indispensable, particularly for the liver which contains intertwined 3D networks enabling the flow of fluids, the sinusoids for blood flow and the BC for bile secretion and flux^9^. Here, we used high resolution multiphoton microscopy and 3D digital reconstructions to generate a comparative dataset of structural changes of human liver tissue from NC, HO, STEA and eNASH patients. We identified a set of zonated morphological alterations that correlate with disease progression, such as a characteristic size distribution of LD and nuclear texture homogeneity, that can be used as tissue biomarkers to distinguish between different stages of NAFLD progression. In addition, the 3D digital reconstruction provided the first evidence that BC integrity is disrupted during NAFLD progression, bringing BC integrity and the mechanisms involved in its maintenance and homeostasis (cell polarity, trafficking, bile flow, BAs turnover, etc.) into focus for NAFLD studies. Based on the geometrical and topological information extracted from the BC, we used a computational personalised model to connect the microanatomy of BC with biliary fluid dynamics within the lobule. Our model predicted high bile pressure in the pericentral area and a significant degree of zonated cholestasis in STEA and eNASH patients, a prediction that was validated by the detection of cholestatic biomarkers in serum. Our data show that geometrical models of human tissues coupled to computational modelling is a powerful strategy to describe human physiology and physiopathology.

The spatially-resolved quantitative analysis of the 3D reconstructions of human liver samples revealed a set of unknown morphological features, ranging from the (sub)cellular (nuclear texture, LD content, polarity) to the tissue level (BC integrity), that are perturbed during NAFLD progression. First, we detected changes in nuclear texture, which have been reported in several diseases^24,25,49^, but have not yet been studied in NAFLD. Indeed, changes in nuclear texture homogeneity in the pericentral hepatocytes may reflect changes in transcriptional activity^27,28^ and could serve as a new component of the histological scores of NAFLD progression. Second, although the accumulation of LD is a characteristic feature of NAFLD, our analysis revealed quantitative changes in their size distribution, with the medium LD mostly present in eNASH and spanning the entire liver lobule. In healthy conditions, the LD number and size are accurately regulated^50^ and changes in LD distribution point at specific alterations in the mechanisms regulating LD biogenesis and catabolism. Third, and most striking, we observed alterations of the apical plasma membrane of hepatocytes and of the BC network. The pericentral hepatocytes showed mislocalization of DPPIV, pointing towards a dysregulation in apical protein trafficking^37^. Interestingly, not all apical proteins were missorted, suggesting that trafficking defects could be pathway-(transcytosis) and/or cargo-specific. Such defects in protein trafficking correlated with the reduction of BC connectivity in the pericentral zone. This is the first time that geometrical and topological properties of the BC are studied in human liver tissue from NAFLD biopsies. The unaltered architecture of the sinusoidal network along the different stages of NAFLD rules out the possibility that this reduction in BC connectivity is simply due to the spatial constrains imposed by the presence of the large pericentral hepatocytes. These results pose the question of how the alterations in apical surface of the hepatocytes collectively result in the compromised connectivity of the BC network.

The altered BC microanatomy and the consequent increase in bile pressure in the pericentral zone suggest that STEA and eNASH livers are affected by a pericentral cholestasis which may contribute to the changes in BA composition observed in NAFLD^42,44,51^. Unimpaired bile flow is essential for normal liver function. Previous studies have documented that bile accumulation, due to its detergent-like properties, can cause liver damage^52,53^ and bile pressure can affect metabolism^54^. The occurrence of zonated cholestasis is a new piece in the NAFLD physiopathology puzzle that contributes to clarify some aspects of the disease so far without explanation, e.g. increase of GGT levels^55^, bile acids in serum^56,57^, upregulation of MRP3 in NASH^58,59^ and the beneficial effect of UDCA treatment in NAFLD^60^, all sign of an ongoing cholestasis^40,61,62^.

In recent years, a lot of research has been devoted to the role of BAs and the activity of their receptor FXR, in NAFLD^42-44^. However, there is currently no explanation for the alterations in BAs composition in blood, the decreased ratio of secondary/primary BAs observed by us (Extended Data Fig. 7) and others^42,44,51^, and whether it correlates with changes in tissue morphology^44^. Our data shed new light on this problem. Our results suggest that the altered BC microanatomy leading to increased bile pressure in the pericentral zone may hamper the ongoing bile acid secretion into BC, as apical pumps (BSEP, MRP2) have to operate against elevated luminal BA concentrations (Extended Data Fig. 9). This could lead to back-flux of primary BA into the blood (Extended Data Fig. 8d), reducing the availability of primary bile acids to be converted into secondary bile acids by the microbiota in the intestine (Extended Data Fig. 7e).

The combination of experimental data with computational models of tissues has proven successful in elucidating pathogenetic mechanisms using animal models^16,63,64^. However, animal models very often fail to mimic human diseases^65^, including NAFLD^66^. In this study, the geometrical models of liver tissue from human biopsies combined with spatially-resolved computational simulations revealed new aspects of NAFLD pathology. This approach may help to identify biomarkers for early disease diagnosis and predict the functional status of the tissue with potential applications in high-definition medicine^1,67,68^.

## Supporting information

3D reconstruction of human liver tissue from NC samples

3D reconstruction of human liver tissue from eNASH samples

Representative pericentral and periportal hepatocytes from NC and eNASH

Representative 3D reconstruction of the bile canaliculi and the sinusoidal networks from the pericentral zone in STEA

## Acknowledgements

We are grateful to Oleksandr Ostrenko, Juan Francisco Miquel Poblete and Sophie Nehring for fruitful discussions, and Sebastian Bundschuh for helping to set up the 2-photon microscope. We thank the Center for Information Services and High Performance Computing (ZIH) of the TU Dresden for the generous provision of computing power. We would also like to thank the following Services and Facilities of the Max Planck Institute of Molecular Cell Biology and Genetics for their support: Light Microscopy Facility (LMF) and the Electron Microscopy Facility.

This work was financially supported by the German Federal Ministry of Education and Research (BMBF) (LiSyM: grant #031L0038 to M.Z., grant #031L0033 to L.B., grant #031L0031 to J.H., DYNAFLOW: grant #031L0082B to M.Z., grant #031L008A to L.B. and SYSBIO II: grant #031L0044 to M.Z.), European Research Council (ERC) (grant #695646 to M.Z.) and the Max Planck Society (MPG).

## Author contributions

F.S-M., J.H. and M.Z. conceived the project. F.S-M., V.M. and S.S. performed the immunofluorescence experiments and imaging. H.M-N. and Y.K. developed the image analysis algorithms. F.S-M., V.M. and H.M-N performed the 3D tissue reconstructions. H.M-N. and F.S-M. performed the data analysis and interpretation of the results. U.R. performed the electron microscopy. A.H., S.H., C.R., and C.S obtained the samples and characterized the patients. D.L. measured bile acids. M.K., F.R., Y.K. and L.B. conceived and developed the mathematical model. M.K and F.R. programmed and simulated the mathematical model and performed statistical analysis. M.K. and L.B. interpreted results and wrote the model description. F.S-M., H.M-N., M.K., Y.K., L.B. and M.Z. wrote the manuscript.

## Competing interests

Authors declare no competing interests

