## Supplementary information for "3D spatially-resolved geometrical and functional models of human liver tissue reveal new aspects of NAFLD progression"

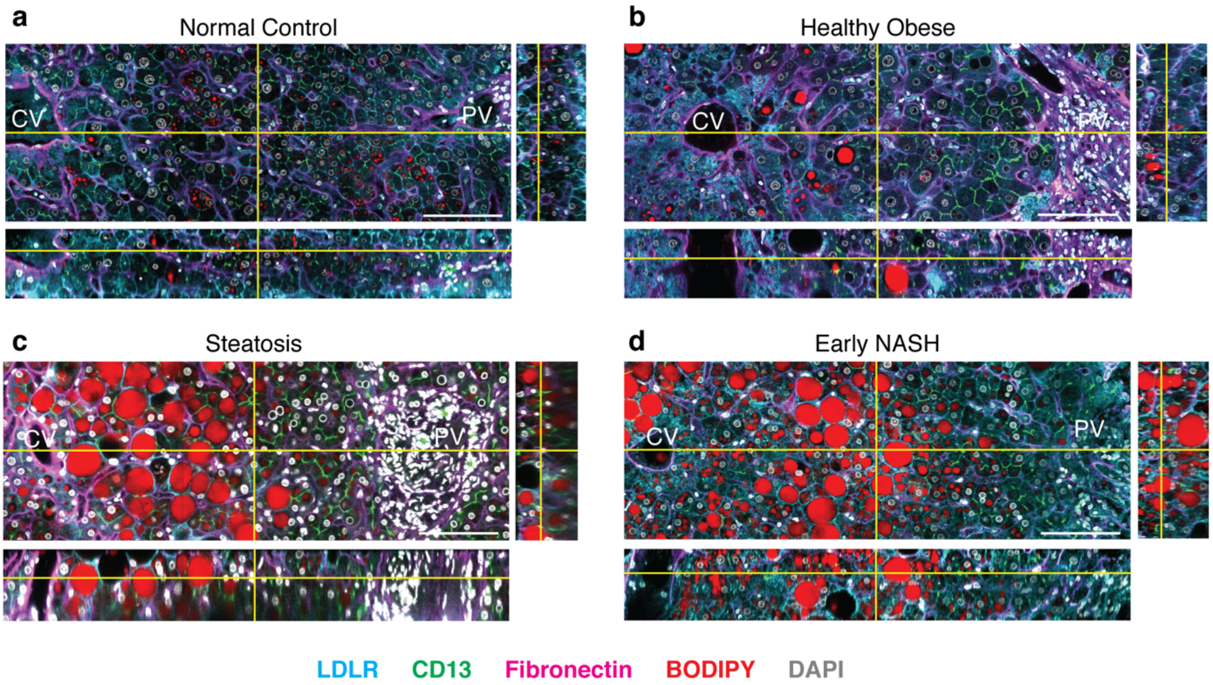


**Extended Data Fig. 1. Immunofluorescence of human liver tissue.** Human liver sections obtained by biopsy (~100 um thick) were stained for bile canaliculi (CD13), sinusoids (fibronectin), nucleus (DAPI), lipid droplets (BODIPY) and cell border (LDLR), optically cleared with SeeDB and imaged at high resolution using multiphoton microscopy (0.3 μm x 0.3 μm x 0.3 μm per voxel). Orthogonal view of NC (**a**), HO (**b**), STEA (**c**) and eNASH (**d**). Scale bar, 50 μm.

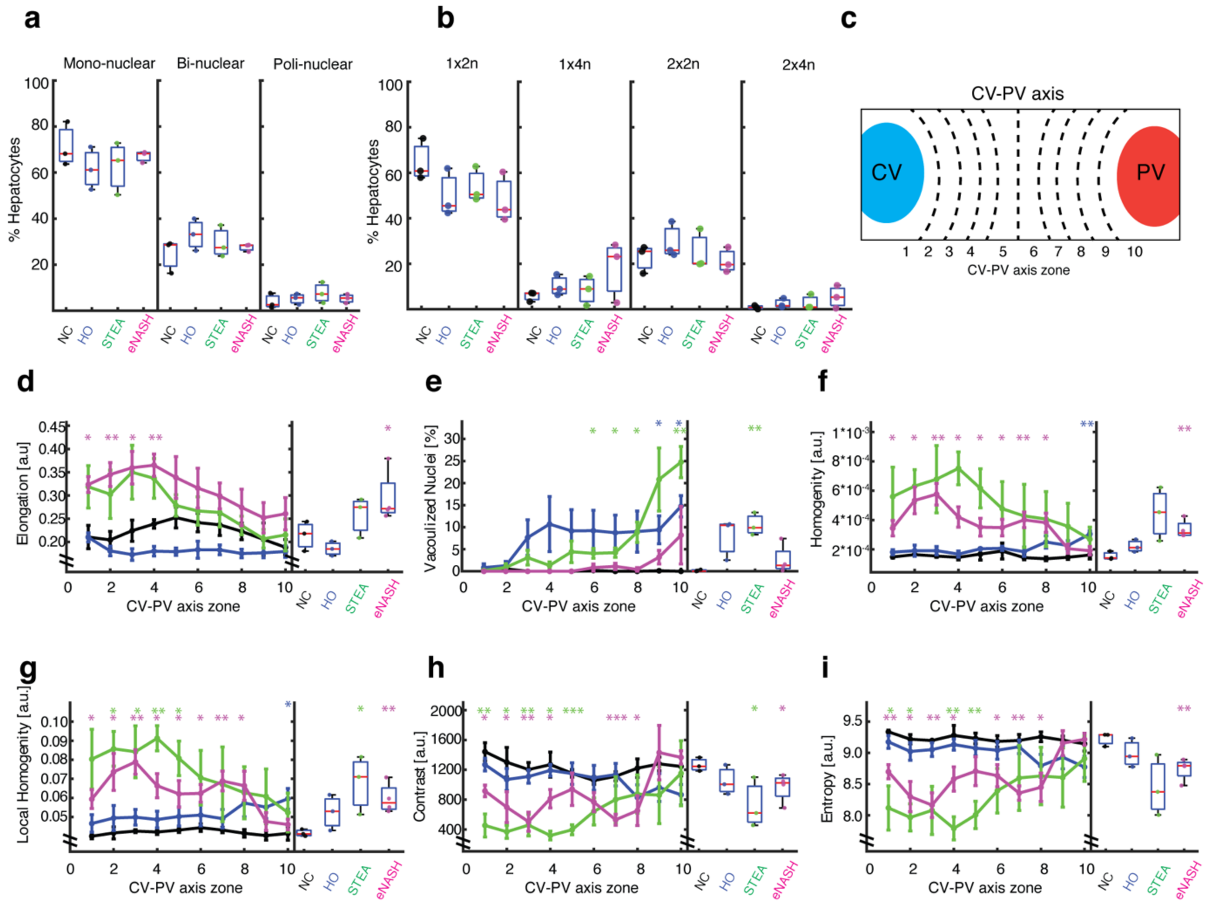


**Extended Data Fig. 2. Morphometric features of the nuclei.** Quantitative characterization of hepatocytes nuclei with respect to the proportion of mono/binuclear cells (**a**) and ploidy (**b**). Only the four major populations (i.e. 1x2n, 1x4n, 2x2n and 2x4n), which account for >90% of the hepatocytes, are shown. **c**, Definition of the zone within the liver lobule. The CV-PV axis was divided in 10 equidistant zones. Zones 1 and 10 are adjacent to the CV and PV, respectively. Quantitative characterization of hepatocytes nuclear elongation (**d**) and texture based on their DAPI intensity (see Methods for details): nuclear vacuolization (**e**), homogeneity (Angular Second Moment) (**f**), local homogeneity (Inverse Difference Moment) (**g**), Contrast (**h**) and Entropy (**i**). NC = 3 samples, HO = 3 samples, STEA = 3 samples, eNASH = 4 samples. Spatially-resolved quantification represented by mean ± SEM per zone and overall quantifications by box-plots. *p-values < 0.05, **p-values < 0.01, ***p-values < 0.001.

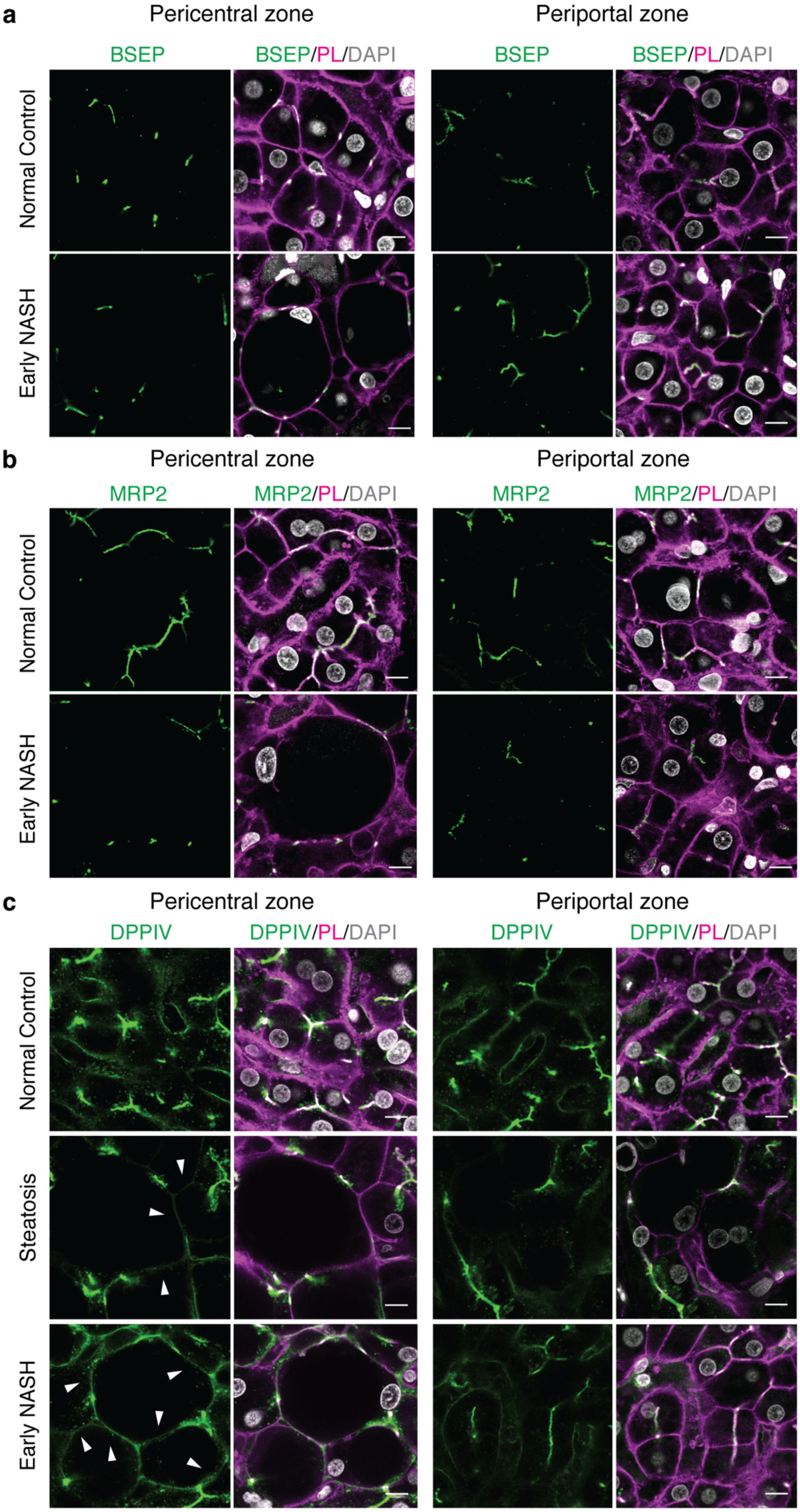

a
Pericentral zone
Periportal zone
BSEP
BSEP/PL/DAPI
Normal Control
Early NASH
b
MRP2
MRP2/PL/DAPI
c
DPPIV
DPPIV/PL/DAPI
Steatosis

**Extended Data Fig. 3. Mislocalization of DPPIV in pericentral hepatocytes in STEA and eNASH.** Representative confocal microscopy images of human liver sections stained for the apical markers BSEP (**a**), MRP2 (**b**) and DPPIV (**c**). Merged images of the apical markers, phalloidin and DAPI are shown in the right panels. Arrowhead indicates the lateral membrane. Scale bar, 10μm.

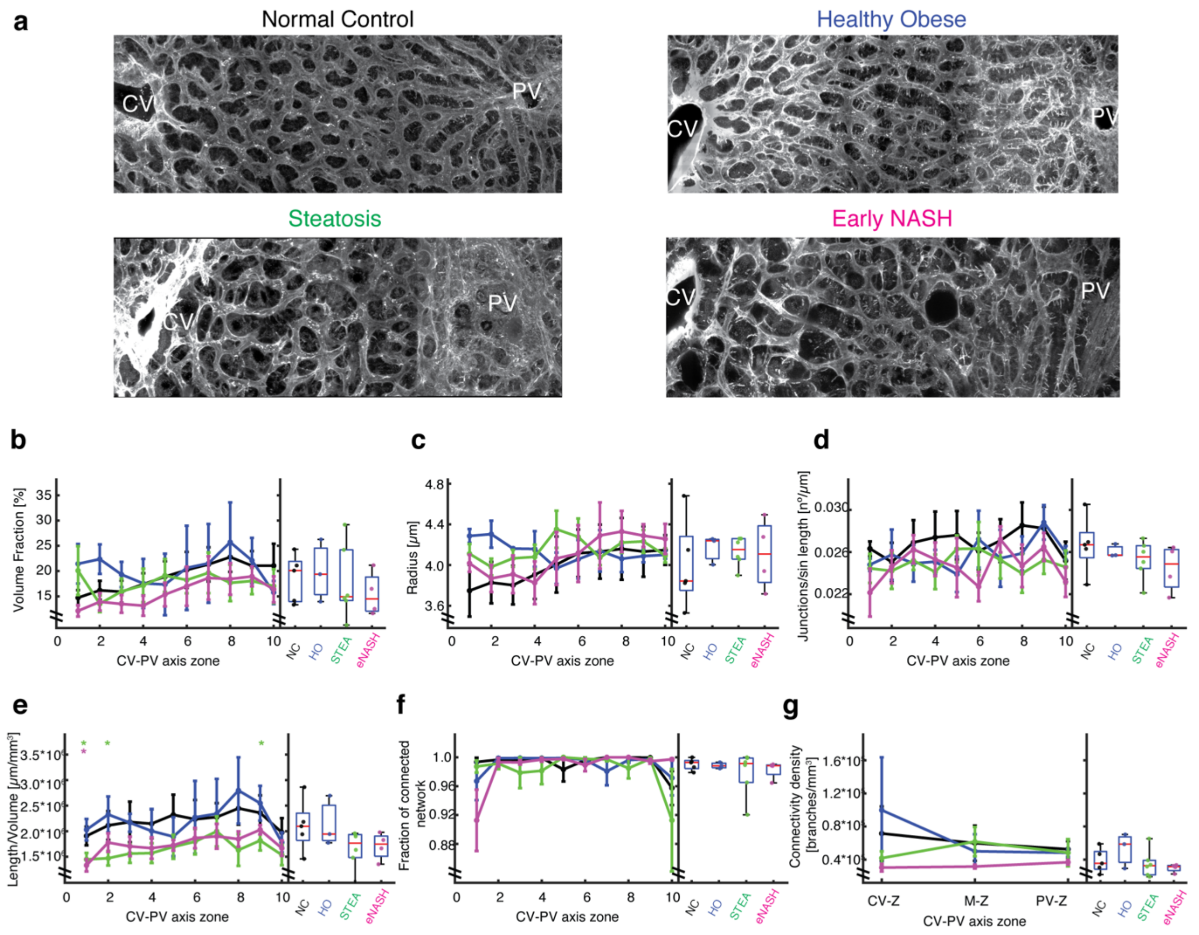


**Extended Data Fig. 4. Structural and topological characterization of the sinusoidal network**. **a**, Representative IF images of fixed human liver tissue sections stained with fibronectin after CAAR. Shown is a maximum projection of a 30 µm z-stack covering an entire CV-PV axis. Quantification of the tissue volume fraction occupied by the sinusoids(**b**), radius (**c**), number of junctions (**d**), total length per unit tissue volume (**e**), fraction of connected network (**f**) and connectivity density (**g**) for the Sinusoidal network along the CV-PV axis and overall. NC = 5 samples, HO = 3 samples, STEA = 6 samples, eNASH = 5 samples. Spatially-resolved quantification represented by mean ± SEM per zone and overall quantifications by box-plots. *p-values < 0.05, **p-values < 0.01, ***p-values < 0.001.

**a**

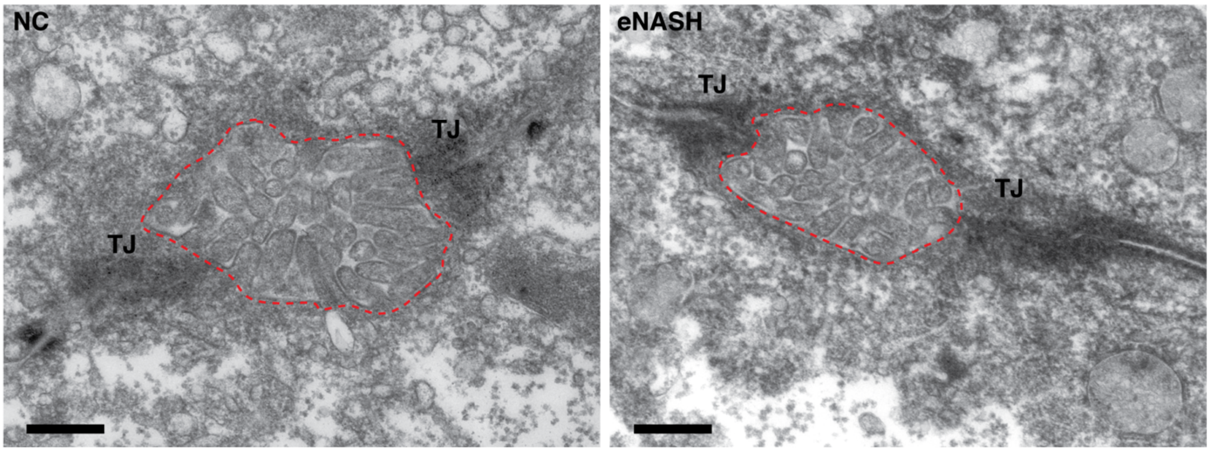


**b**

Percentage of free lumen

| | Pericentral zone | | Periportal zone | |
|---|---|---|---|---|
| | mean | sem | mean | sem |
| NC | 28.8 | 0.32 | 29.5 | 1.38 |
| HO | 30.3 | 0.42 | 29.1 | 0.81 |
| STEA | 24.9 | 1.83 | 27.1 | 1.01 |
| eNASH | 27.0 | 0.84 | 23.8 | 2.76 |

**Extended Data Fig. 5. Estimates for a fraction of free lumen in total volume of a bile canaliculus. a**, Representative images of bile canaliculi for NC and eNASH liver tissue samples, used for making the estimates. Microvilli are well preserved. A red dashed line indicates lumen of a bile canaliculus. TJ, tight junction. Scalebar, 500 nm. **b**, Estimation of fraction of free lumen by stereological point counting (the Cavalieri estimator). For each set of samples and each region (central / portal vein) a minimum of five EM images was used. NC = 3 samples, HO = 3 samples, STEA = 3 samples, eNASH = 3 samples, mean ± SEM.

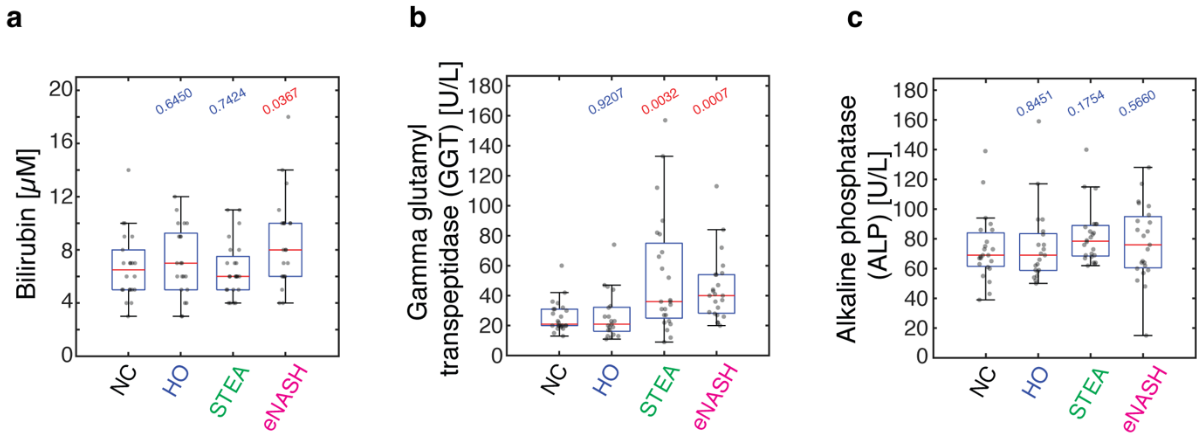


**Extended Data Fig. 6. Increased serum cholestatic markers during disease progression.** The levels of bilirubin (**a**), GGT (**b**) and ALP (**c**) were measured in the serum of the patients and represented in box-plots. NC = 25 samples, HO = 25 samples, STEA = 24 samples, eNASH = 26 samples. Two-sided t-test assuming unequal variances. P-values are indicated in the plots.

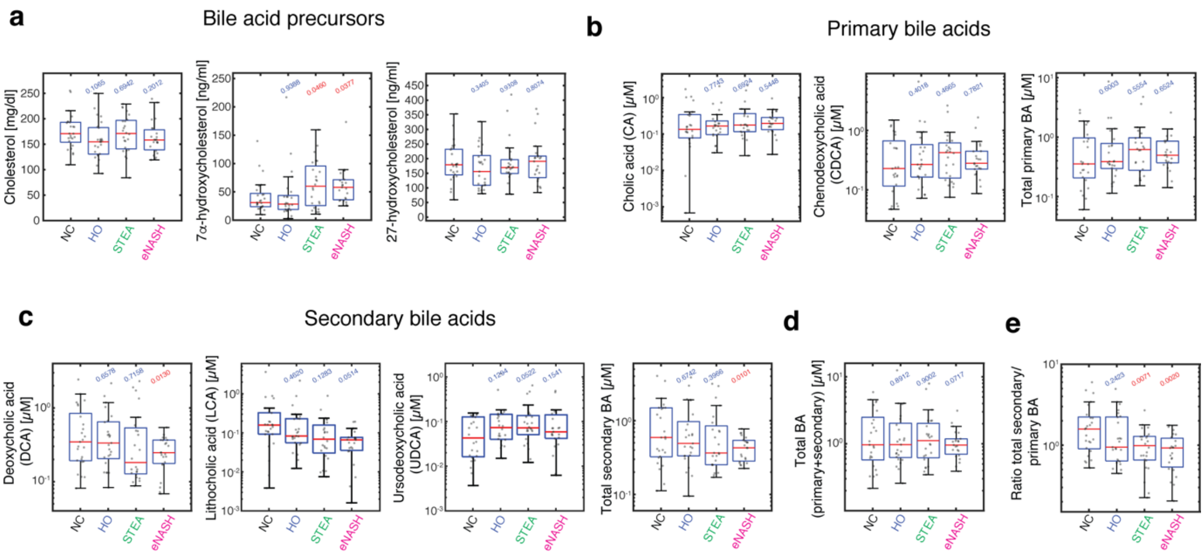


**Extended Data Fig. 7. Profile of serum bile acids during disease progression.** Individual BAs and their precursors were measured in the serum of the patients and represented in box-plots. **a**, BA precursors (cholesterol, 7α-hydroxycholesterol and 27-hydroxycholesterol). **b**, Individual and total primary BAs (CA, CDCA). **c**, Individual and total secondary BAs (DCA, LCA, UDCA). **d**, Total BAs. **e**, Ratio secondary to primary BAs. NC = 25 samples, HO = 25 samples, STEA = 24 samples, eNASH = 26 samples. Two-sided t-test assuming unequal variances. P-values are indicated in the plots.

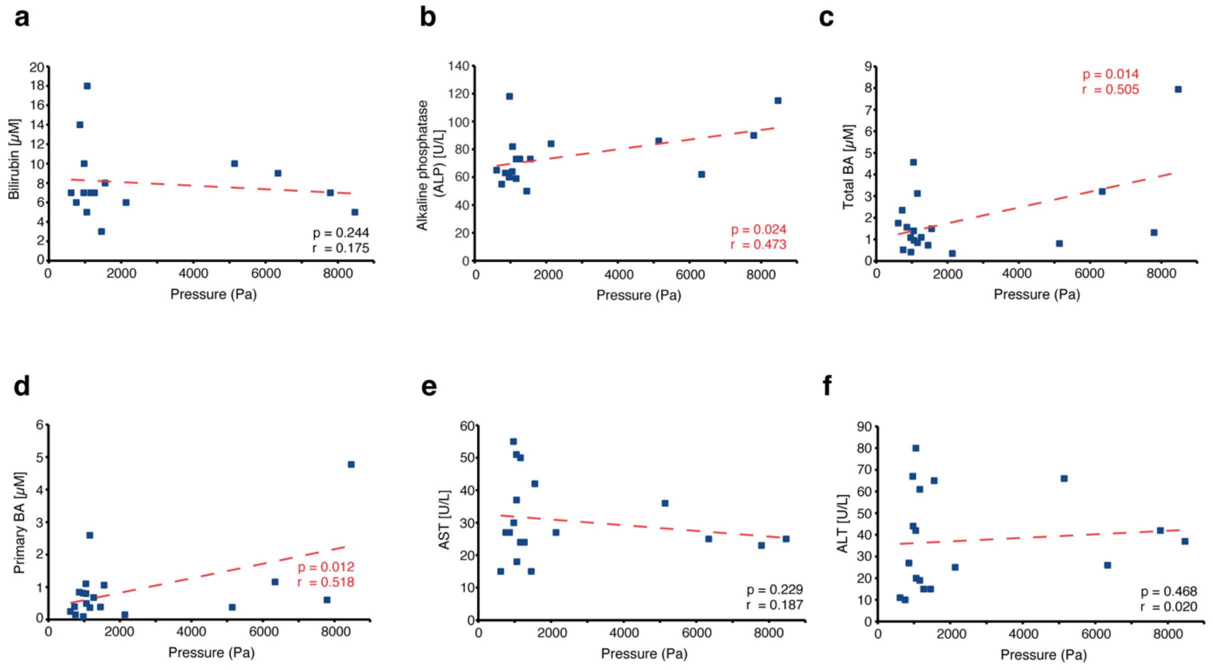


**Extended Data Fig. 8 Scatter plots and regression analysis of measured liver enzymes and bile acids.** (**a**) bilirubin, (**b**) ALP, (**c**) total BAs, (**d**) primary BAs, (**e**) secondary BAs, (**f**) AST and (**g**) ALT measured in the serum versus the model-derived pericentral pressure in individual patients from all groups. One-sided t-test. P-values and Pearson correlation coefficient are indicated in the plots.

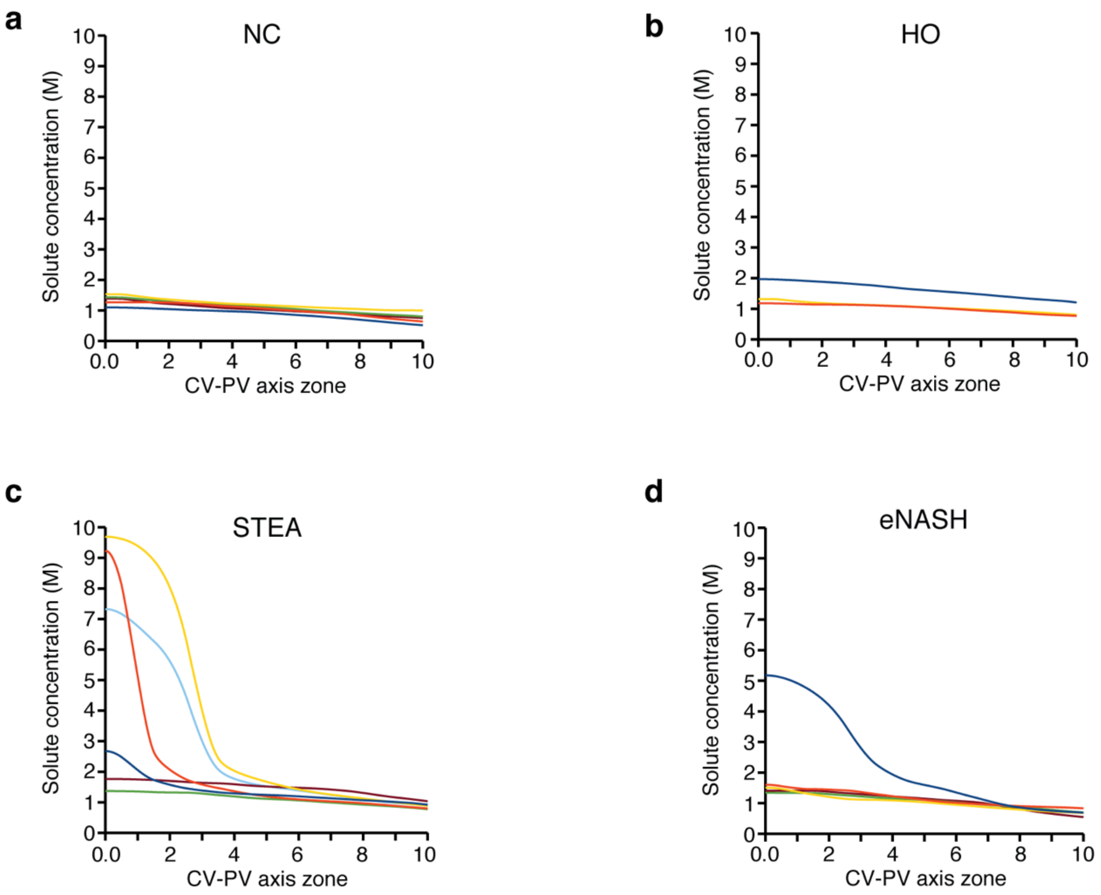


**Extended Data Fig. 9 Model-derived concentration profiles for each patient in each patient group.** (**a**) NC, (**b**) HO, (**c**) STEA) and (**d**) eNASH. NC = 5 samples, HO = 3 samples, STEA = 6 samples, eNASH = 5 samples.

| | | Demographic | | | | | | Histology | | | | | | Blood Test | | | | |
|---|---|---|---|---|---|---|---|---|---|---|---|---|---|---|---|---|---|---|
| **ID** | **Group** | **Sex** | **Age (years)** | **Weight (kg)** | **BMI** | **Diabetes Typ II** | **Surgery indication** | **NAS Score** | **NAS Fat** | **NAS Balloning** | **NAS Inflamation** | **% fat** | **Fibrosis** | **GGT (U/l)** | **AP (U/l)** | **total bilirubin (mg/dl)** | **ALT (U/l)** | **AST (U/l)** |
| 6724 | NC | female | 92 | 48 | 20.5 | yes | pancreatic cancer | 0 | 0 | 0 | 0 | 0 | no | 60 | 87 | 0.650 | 42 | 51 |
| 6929 | NC | female | 84 | 60 | 24.65 | no | colorectal cancer | 0 | 0 | 0 | 0 | 3 | no | 20 | 73 | 0.420 | 15 | 24 |
| 6936 | NC | male | 75 | 75 | 25.35 | no | colorectal cancer | 1 | 1 | 0 | 0 | 5 | no | 20 | 65 | NA | 11 | 15 |
| 7012 | NC | female | 54 | 65 | 21.97 | no | gastric cancer | 0 | 0 | 0 | 0 | 0 | no | 16 | 35 | 0.300 | 12 | 22 |
| 7173 | NC | female | 54 | 77 | 26.03 | no | gastric cancer | 0 | 0 | 0 | 0 | 0 | no | 42 | 118 | 0.585 | 44 | 30 |
| 7194 | NC | male | 62 | 84 | 27.12 | no | pancreatic cancer | 0 | 0 | 0 | 0 | 0 | no | 35 | 63 | 0.819 | 27 | 27 |
| 6758 | HO | female | 36 | 110 | 40.9 | no | sleeve gastrectomy | 1 | 1 | 0 | 0 | 5 | no | 19 | 50 | 0.190 | 15 | 16 |
| 6922 | HO | male | 68 | 95 | 31.7 | yes | colorectal cancer | 1 | 1 | 0 | 0 | 5 | no | 31 | 67 | 0.250 | 33 | 20 |
| 7230 | HO | female | 36 | 126 | 40.22 | no | gastric bypass | 0 | 0 | 0 | 0 | 0 | no | 17 | 55 | 0.351 | 10 | 27 |
| 6922 | HO | male | 68 | 95 | 31.7 | yes | colorectal cancer | 1 | 1 | 0 | 0 | 5 | no | 31 | 67 | 0.250 | 33 | 20 |
| 6610 | STEA | male | 41 | 185 | 59.7 | yes | gastric bypass | 3 | 2 | 1 | 0 | 50 | no | 81 | 90 | 0.430 | 42 | 23 |
| 6718 | STEA | female | 54 | 90 | 31.9 | no | colorectal cancer | 0 | 0 | 0 | 0 | 3 | no | 157 | 115 | 0.310 | 37 | 25 |
| 6757 | STEA | male | 33 | 160 | 46.7 | no | sleeve gastrectomy | 2 | 2 | 0 | 0 | 50 | no | 36 | 64 | 0.280 | 80 | 37 |
| 6503 | STEA | female | 41 | 198 | 54.3 | no | sleeve gastrectomy | 1 | 1 | 0 | 0 | 25 | no | 21 | 84 | 0.360 | 25 | 27 |
| 7137 | STEA | male | 67 | 115 | 33.24 | yes | gastric cancer | 3 | 3 | 0 | 0 | 90 | yes | 133 | 62 | 0.526 | 26 | 25 |
| 6921 | eNASH | male | 55 | 169 | 75.11 | no | gastric bypass | 4 | 2 | 1 | 1 | 40 | no | 54 | 86 | 0.560 | 66 | 36 |
| 6980 | eNASH | female | 51 | 150 | 51.00 | no | gastric bypass | 4 | 2 | 1 | 1 | 60 | no | 37 | 82 | 1.040 | 20.2 | 18.5 |
| 7041 | eNASH | female | 44 | 155 | 50.61 | no | gastric bypass | 2 | 1 | 0 | 1 | 40 | no | 29 | 96 | 0.585 | 31 | 28 |
| 7157 | eNASH | male | 44 | 150 | 40.27 | yes | gastric bypass | 4 | 2 | 1 | 1 | 40 | yes | 44 | 65 | 0.585 | 60 | 35 |
| 7188 | eNASH | female | 51 | 114 | 45.09 | no | gastric bypass | 3 | 2 | 0 | 1 | 40 | no | 28 | 59 | 0.409 | 19 | 24 |
| 7251 | eNASH | male | 34 | 163 | 48.15 | no | sleeve gastrectomy | 3 | 3 | 0 | 0 | 70 | no | 40 | 60 | 0.409 | 67 | 55 |
| 7344 | eNASH | female | 58 | 155 | 51.20 | yes | gastric bypass | 3 | 2 | 0 | 1 | 50 | no | 113 | 73 | 0.468 | 65 | 42 |

**Supplementary Table 1:** Patient characteristics by group.

| Primary Abs | Dilution | Source | Identifier | With CAAR | Without CAAR | Description |
|---|---|---|---|---|---|---|
| Rat polyclonal antibody against CD13 | 1/500 | Acris Antibodies | Cat# SM2298P | - | - | Canaliculi marker |
| Mouse monoclonal antibody against CD13 | 1/50 | Santa Cruz | Cat# sc-136484 | ++++ | - | Canaliculi marker |
| Rabbit monoclonal antibody against CD13 | 1/100 | Abcam | Cat# ab108310 | - | - | Canaliculi marker |
| Goat polyclonal antibody against DPPIV | 1/100 | R&D systems | Cat# AF1180 | +++ | ++++ | Canaliculi marker |
| Goat polyclonal antibody against DPPIV | 1/200 | R&D systems | Cat# AF954 | - | - | Canaliculi marker |
| Mouse monoclonal antibody against ZO-1 | 1/500 | Thermo Fisher Scientific | Cat# 33-9100 | - | - | Canaliculi marker |
| Rabbit polyclonal antibody against BSEP | 1/2000 | Sigma-Adrich | Cat# HPA019035 | ++ | +++ | Canaliculi marker |
| Mouse monoclonal antibody against BSEP | 1/200 | Santa Cruz | Cat# 33-9100 | - | - | Canaliculi marker |
| Mouse monoclonal antobody against MRP2 | 1/100 | Abcam | Cat# ab3373 | ++ | ++ | Canaliculi marker |
| Rabbit polyclonal antibody against fibronectin | 1/1000 | LifeSpan BioScience | Cat# LS-B2318 | ++++ | +++ | Sinusoidal marker |
| Rabbit polyclonal antibody against fibronectin | 1/1000 | Millipore | Cat# AB2033 | - | - | Sinusoidal marker |
| Rabbit polyclonal antibody against colagen type-I | 1/100 | Millipore | Cat# AB765P | - | - | Sinusoidal marker |
| Rabbit polyclonal antibody against laminin | 1/5000 | Sigma-Adrich | Cat# L9393 | - | - | Sinusoidal marker |
| Rabbit polyclonal antibody against laminin | 1/1000 | LifeSpan BioScience | Cat# LS-B4754 | - | - | Sinusoidal marker |
| Goat polyclonal antibody against ASGPR1 | 1/200 | Santa Cruz | Cat# sc-13469 | - | - | Sinusoidal marker |
| Goat polyclonal antibody against Flk-1 | 1/200 | R&D systems | Cat# AF644 | ++ | ++ | Sinusoidal marker |
| Rat monoclonal antibody against CD31 | 1/200 | eBioscience | Cat# 14-0311 | - | - | Sinusoidal marker |
| Rat monoclonal antibody against CD29 | 1/50 | BD Biosciences | Cat# 550531 | - | - | Sinusoidal marker |
| Rat monoclonal antibody against CD16/32 | 1/500 | BD Biosciences | Cat# 553142 | - | - | Sinusoidal marker |
| Rat monoclonal antibody against CD144 | 1/500 | eBioscience | Cat# 14-1441 | - | - | Sinusoidal marker |
| Goat polyclonal antibody against Neuropilin-1 | 1/200 | R&D systems | Cat# AF566 | - | - | Sinusoidal marker |
| Goat polyclonal antibody against CD14 | 1/100 | Novus | Cat# NB100-2807 | +++ | ++ | Sinusoidal marker |
| Chicken polyclonal antibody against LDLR | 1/150 | Sigma-Adrich | Cat# SAB3500286 | ++++ | ++ | Cell border |
| Mouse monoclonal antibody agaist E-Cadherin | 1/200 | BD Biosciences | Cat# 610182 | +++ | +++ | Cell border |
| Rabbit mnoclonal antibody against E-Cadherin | 1/200 | Cell Signaling | Cat# 3195 | ++ | - | Cell border |
| Rat monoclonal antibody against Integrin β1 | 1/200 | Millipore | Cat# 1997 | - | - | Cell border |

| Secondary Abs and dyes | Dilution | Source | Identifier | With CAAR | Without CAAR | Description |
|---|---|---|---|---|---|---|
| Donkey anti mouse Alexa Fluor 555 | 1/1000 | Invitrogen | Cat# A31570 | ++++ | ++++ | |
| Donkey anti rabbit Alexa Fluor 555 | 1/1000 | Invitrogen | Cat# A31572 | ++++ | ++++ | |
| Donkey anti goat Alexa Fluor 568 | 1/1000 | Invitrogen | Cat# A11057 | ++++ | ++++ | |
| Donkey anti goat Alexa Fluor 594 | 1/1000 | Invitrogen | Cat# A11058 | ++++ | ++++ | |
| Donkey anti rabbit Alexa Fluor 594 | 1/1000 | Invitrogen | Cat# A21207 | ++++ | ++++ | |
| Donkey anti mouse Alexa Fluor 647 | 1/1000 | Invitrogen | Cat# A31571 | ++++ | ++++ | |
| Donkey anti rabbit Alexa Fluor 647 | 1/1000 | Invitrogen | Cat# A31573 | ++++ | ++++ | |
| Donkey anti chicken Alexa Fluor 647 | 1/25 | Millipore | Cat# AP194SA6 | ++++ | ++++ | |
| Phalloidin Alexa Fluor 647 | 1/100 | Invitrogen | Cat# A22287 | - | ++++ | Cell border |
| Phalloidin Alexa Fluor 488 | 1/100 | Invitrogen | Cat# A12379 | - | ++++ | Cell border |
| BODIPY 493/503 | 1/10000 | Invitrogen | Cat# D3922 | ++++ | ++++ | Lipid Droplets |
| DAPI 1mg/ml | 1/1000 | Sigma-Adrich | Cat# D8417 | ++++ | ++++ | Nuclei |

**Supplementary Table 2:** Antibodies and dyes tested for staining of human liver tissue.

**Supplementary Video 1:** 3D reconstruction of human liver tissue from NC samples. Central vein (light blue), portal vein (orange), bile canaliculus (green), sinusoids (magenta), lipid droplets (red), nuclei (random colours) and hepatocytes (random colours).

**Supplementary Video 2:** 3D reconstruction of human liver tissue from eNASH samples. Central vein (light blue), portal vein (orange), bile canaliculus (green), sinusoids (magenta), lipid droplets (red), nuclei (random colours) and hepatocytes (random colours).

**Supplementary Video 3:** Representative pericentral and periportal hepatocytes from NC and eNASH liver tissue samples. Apical (green), basal (magenta) and lateral (grey) plasma membrane domains, nuclei (random grey shades) and lipid droplets (red).

**Supplementary Video 4:** Representative 3D reconstruction of the bile canaliculi (green) and the sinusoidal (magenta) networks from the pericentral zone in STEA.

### Methods

#### Human liver samples

Liver samples were obtained intraoperatively in patients in whom an intraoperative liver biopsy was indicated on clinical grounds such as exclusion of liver malignancy during major oncologic surgery or assessment of liver histology during bariatric surgery. Standardized histopathologic assessment was performed by a single pathologist based on the NAFLD activity score (NAS)[1]. The samples were divided into 4 groups: normal control (NC), healthy obese (HO), steatosis (STEA) and early NASH (eNASH). None of the individuals underwent preoperative chemotherapy and liver histology demonstrated absence of both cirrhosis and malignancy. Biopsy specimens were fixed immediately in 4% paraformaldehyde 0.1%Tween-20/PBS for 2-3 days. Patients with evidence of viral hepatitis, hemochromatosis, or alcohol consumption greater than 20 g/day for women and 30 g/day for men were excluded. All patients provided written informed consent. The study protocol was approved by the institutional review board (Ethikkommission der Medizinischen Fakultät der Universität Kiel, D425/07, A111/99) before study commencement.

#### Immunolabeling, optical clearing and imaging

For the immunostainings where heat-induced epitope retrieval was required, liver slices were transferred to an eppendorf tube with citrate buffer pH 6.0 (Sigma-Aldrich Cat# C9999) and heated for 30 min at 80ºC. Next, immunolabeling (Supplementary Table 02) and optical clearing (modified version of SeeDB[2]) were performed as described previously[3].

Liver samples were imaged (0.3 µm voxel size) in an inverted multiphoton laser-scanning microscope (Zeiss LSM 780 NLO) using a 63x 1.3 numerical aperture glycerol immersion objective (Zeiss). DAPI and phalloidin A647 were excited at 790nm using a Chameleon Ti-Sapphire 2-photon laser and detected with non-descanned detectors (NDD). Alexa Fluor 488, 555 and 594 were excited with 488, 561 and 594 laser lines and detected with Gallium arsenide phosphide (GaAsp) detectors.

#### 3D tissue reconstruction

The different components of live tissue (i.e. BC and sinusoidal networks, nuclei, lipid droplets and hepatocytes) were reconstructed from high-resolution (voxel size: 0.3 x 0.3 x 0.3 µm) fluorescent image stacks (~100 µm depth) of fixed liver tissue stained by specific antibodies and/or small fluorescent molecules (i.e. CD13, fibronectin, DAPI, BODIPY, and LDLR). To cover entire CV-PV axes, tiles of 2x2 or 3x1 image stacks were stitched using the Image Stitching plug-in of Fiji[4]. Then, all images were reconstructed using the software MotionTracking as described in[5]. Briefly, all stitched images were desoised using the Bayesian foreground/background discrimination (BFBD) de-noising algorithm proposed in[5]. Next, the tubular structures (BC and sinusoidal networks) as well as nuclei were segmented using Maximum entropy local thresholding algorithm. Artefacts generated by the segmentation (holes and tiny isolated object) by standard morphological operations (opening/closing). The triangulation mesh of the segmented surfaces was generated by the cube marching algorithm and tuned using an active mesh approach. In the case of tubular structures, representations of the skeleton, refereed as central lines, of the networks were generated. Central lines are represented as 3D graphs. Finally, the shape of the cell surface (i.e. hepatocytes) was found using an active mash expansion from the reconstructed nuclei. For details, refer to[5].

**Morphological spatial analysis**

To account for the variability of different morphological parameters along the liver lobule, the CV-PV axis was computationally divided in 10 equidistance zone. The zones were defined in terms of the distances to the closest CV and PV, as follows:

$$zone_i = \left\lfloor N \times \frac{d_{cv}}{d_{cv} + d_{pv}} \right\rfloor + 1$$

For $i = 1{:}N$, where $N = 10$, is the number of zones, $\lfloor \quad \rfloor$ is the floor function, $d_{cv}$ and $d_{pv}$ are the distances to the closest CV and PV respectively. The average value of the different morphological parameters was calculated in every zone.

**Nucleus vacuolization**

To determine if a nucleus is vacuolated, we calculated the DAPI mean intensity in the middle of the nucleus (inside a sphere of 1.2 µm located at the centre of the nucleus) as well as in the inner surface of it (over a layer of thickness 0.6 µm). If the inner DAPI intensity was 3 times lower than the DAPI intensity in the surface, the nucleus was defined as vacuolated.

**Nucleus homogeneity**

We analysed the nuclear texture based on the Haralick texture features[6] measured over images of DAPI. We examined the four features related to the texture homogeneity/heterogeneity of the image, i.e. homogeneity (Angular Second Moment), local homogeneity (Inverse Difference Moment), Contrast and Entropy. To avoid boundary artefacts, we analysed only the intensity of DAPI inside a cube located in the centre of the nucleus and of length equal to half of the nucleus radius. All features were extracted from the normalized grey-level co-occurrence matrix (256 grey-levels) averaged over the 13 directional co-occurrence matrices (in 3D) at distances from 1 to 3 pixels. Vacuolated nuclei were excluded from the analysis.

**Network connectivity**

Two approaches were used to quantify the connectivity of the tubular networks (i.e. BC and sinusoidal networks). The first one, also referred as 'fraction of connected network', is based on the Central lines of the network and defined as the ratio between the length of the largest connected graph of the network and the total length of the network. The second one is based on the segmented 3D image of network and calculates the connectivity density using the Euler characteristic of the network (maximum number of branches that can be removed before the network is separated in two parts)[7,8]. The former calculation was performed in Fiji[9] using the plugin BoneJ[10].

**Electron microscopy**

For the ultrastructural analysis of bile canaliculi, we used 100-µm thick vibratome sections of human liver biopsy tissue originally fixed with 4% PFA for several days and then stored in PBS. Before embedding, vibratome sections were fixed with 1% glutaraldehyde in 200 mM HEPES at least overnight and then cut into small pieces. Next, tissue was post-fixed with 1% osmium tetroxide prepared in 1.5% potassium ferricyanide, for 1 hr at room temperature. After washing with water, tissue was incubated with 1% tannic acid dissolved in 100 mM HEPES, pH 7.0 for 20 min, followed by incubation with 1% disodium sulfate for 5 min, and then by several washes with water. After that tissue was incubated with 2% aqueous uranyl acetate for 2 hrs at room temperature and protected from light. A graded ethanol series was used for dehydration: 70-80-90-96%, each 10 min, followed by absolute ethanol, 4x 15 min. Tissue was progressively infiltrated with epon over 24 hrs. Finally, tissue pieces were flat embedded between two teflon-coated glass slides and heat polymerized overnight. Embedded tissue pieces were remounted for longitudinal sectioning. Periportal or pericentral regions were selected on 1-µm sections, stained with methylene blue-azur II and examined in a light microsope. Then 70-80-nm thin sections were cut. These were stained with 0.4% lead citrate for 1 min and imaged in the Tecnai T12 transmission electron microscope (ThermoFisher), equipped with an axial Tietz CCD camera (TVIPS). Images of bile canaliculi were taken at 6,800x maginification by systematic and random screening. Bile canaliculi with poorly preserved microvilli were not included in the analysis.

To estimate a fraction of free lumen (i.e., lumen not occupied by microvilli) in bile canaliculi, we applied stereological point counting (the Cavalieri estimator), using Fiji[9] software. A test grid of vertical and horizontal lines (area per point 5000 nm$^2$) was laid over images. Cross points over total lumen profile area and over free lumen profile area were separately counted. For each set of samples and each region (central / portal vein) a minimum of five EM images was used, and counts obtained from individual images were summed up to obtain total counts. The analysis was performed in a blinded way.

**Bile flow model in the hepatic lobule**

We consider a three-dimensional domain $\Omega$ with an osmolyte influx density $g(\vec{x})$ and a fluid influx density $j(\vec{x})$, leading to a lumped concentration profile $c(\vec{x})$ for osmolytes throughout the bulk of the domain. The bulk velocity field of the fluid is given in most generality by Darcy's law according to which velocity $v(\vec{x})$ is proportional, with a resistance coefficient $-K(\vec{x})$, to the pressure gradient $\nabla p(\vec{x})$. We assume that diffusion of osmolytes is negligible on the length scale of paths through the BC network, hence also in our bulk model. At steady-state, this system is described by conservation of mass for the osmolytes as well as the fluid and the connection between pressure gradient and velocity as follows:

$$\nabla\vec{v}(\vec{x}) = j(\vec{x}), \qquad \nabla(c(\vec{x})\vec{v}(\vec{x})) = g(\vec{x}), \qquad \nabla p(\vec{x}) = -K(\vec{x})\vec{v}(\vec{x}) \ .$$

To first approximation, the liver lobule can be assumed to have cylindrical symmetry. Therefore, we take $\Omega$ as a cylindrical domain with radial symmetry and radius $L$, approximating a single prismatic liver lobule. Hence, we need to consider only the radial variable which will be denoted by $\rho$. Following [11,12], we consider the fluid influx $j$ as proportional to the difference in osmotic pressure ($RTc$) and hydrostatic pressure. Using the divergence in cylinder coordinates $\nabla\vec{v} = (\rho w)'/\rho$, with the radial velocity component $w$ and the apostrophe denoting the derivative with respect to $\rho$, we obtain the following equations:

$$(\rho w)'/\rho = \kappa A(\rho)[RTc - p]$$
$$(\rho c w)'/\rho = g(\rho)$$
$$p' = -K(\rho)w$$

where $A(\rho)$ denotes the spatial density of hepatocyte apical membrane as osmotically active surface and $\kappa$ denotes the membrane permeability for water. The second equation can be readily integrated as follows

$$\rho c w = \int_0^\rho \tilde{\rho} \ \ g(\tilde{\rho} \ \ )d\tilde{\rho} \qquad .$$

The integral expression represents the cumulative osmolyte source flux inside radius $\rho$ and will be abbreviated as $G(\rho)$. Using this equation, we can eliminate the concentration term in the equation for $w'$ as follows:

$$(\rho w)'/\rho = \kappa A[RT\, G/(\rho w) - p]$$

or rewritten

$$w' = \kappa A[RTG/(\rho w) - p] - w/\rho \quad .$$

Together with the equation for $p'$, these are coupled first-order ordinary differential equations (ODEs) with the mixed boundary conditions $w(\rho_0) = 0$ and $p(L) = p_L$ where $\rho_0$ is the radius of the central vein and $p_L$ is the hydrostatic pressure in the bile duct. Since some model parameters such as $A(\rho)$ and $K(\rho)$ will below be derived directly from the quantification of image data, these parameters are given as heterogeneous profiles and the ODEs therefore possess heterogeneous coefficients.

We can nondimensionalize $\rho$ as follows:

$$\bar{\rho} = \rho/L \quad ,$$

which modifies the equations as follows (immediately dropping the bar):

$$w' = \kappa AL[RTLG/(\rho w) - p] - w/\rho$$

$$p' = -LKw \quad .$$

Note that there is no singularity at $\rho = \rho_0$ even though $w$ appears in the denominator because we have, after applying l'Hospital's rule:

$$\lim_{\rho\to\rho_0} \frac{G}{w} = \frac{\rho_0 g(\rho_0)}{w'(\rho_0)}$$

and hence a finite limit for $\rho \to \rho_0$.

The equations are solved numerically using the shooting method[13]. For this method, we set the initial values to $w(\rho_0) = 0$ (the proper condition) and $p(\rho_0) = p^*$ with some arbitrarily chosen value $p^*$. We solve the ODEs for $w$ and $p$ with an explicit integration scheme till $\rho = 1$. Next, we iteratively updated $p^*$ until $p(1) = p_L$. To avoid any division by $w = 0$ at $\rho = \rho_0$ in the first integration step, we made use of the above equation for the limit $\rho \to \rho_0$. This resulted in an algebraic equation for $w'(\rho_0)$:

$$w'(\rho_0) = \frac{\kappa AL^2 RTg(\rho_0)}{w'(\rho_0)} - \kappa ALp(\rho_0)$$

which has the solution

$$w'(\rho_0) = -s/2 + \sqrt{s^2/4 + t}$$

with

$$t = \kappa AL^2 RT\, g(\rho_0) \quad \text{and} \quad s = \kappa ALp(\rho_0) \quad .$$

The equations were integrated using operator splitting: the explicit solution was used for the $w/\rho$ term, and an Euler method was used for the remaining terms.

For the determination of $K$, we considered a network of tubes with radii $r(\rho)$ and tortuosity $\tau$ that cause a bulk porosity of $\varepsilon(\rho)$. The tortuosity of a curve is defined as the ratio between its length and the distance between its end points. It is determined from individual patient data ($\tau$ ranges from 1.5 to 2.3). The porosity $\varepsilon(\rho)$ is determined from patient data as the product of the BC volume fraction  and the BC connectivity profile, amounting to pruning of branches that do not relay the flow. Where BC connectivity was quantified as 0, we set it to 0.01 assuming that at least some remote connection may exist and thereby potentially underestimating the pressure profile. We take into account that a significant portion of the BC lumen is taken up by microvilli. From our EM measurements, we know the fraction of free to total lumen $\alpha(\rho)$, therefore the effective porosity and radius are given by $\varepsilon_{\mathrm{BC}}(\rho) = \alpha(\rho)\,\varepsilon(\rho)$ and $r_{\mathrm{BC}}(\rho) = \sqrt{\alpha(\rho)}\; r(\rho)$. With these quantities $K(\rho)$ is given exclusively in terms of a patient's data as[14]

$$K(\rho) = \frac{8\mu\tau}{\varepsilon_{\mathrm{BC}}(\rho) r_{\mathrm{BC}}(\rho)^2} \quad .$$

We use the universal gas constant $R$ = 8.314 J/mol/K and $T$ = 293 K. The values for $A(\rho)$ and $L$ are of the order of 5x10$^{4}$x 1/m and 4x10$^{-4}$ m, respectively, but the exact values are taken from our measurements for each individual patient sample. The membrane permeability $\kappa$ = 3x10$^{-10}$ m/(sPa) had been estimated previously. We consider an outer pressure $p_L = 100\ \mathrm{Pa}$ (estimated according to measurements reported in[15]). The central vein radius was measured to be 2x10$^{-5}$ m in our samples.

We determine the approximate osmolyte flux $g$ as follows: about 500 ml of bile with an osmolarity of 300 mmol/l drain every day from a human liver of 1 l in size. This means that 150 mmol of osmolytes are secreted through the apical membranes of hepatocytes every day. This leads to $g$ = 1.7x10$^{-3}$ mol/m$^3$/s. The viscosity of bile was chosen as $\mu$ = 8x10$^{-4}$ Pa s which is close to the viscosity of water[16].

Therewith all parameter values have been determined from previously reported data and geometrical measurements in this work such that no parameter fitting is necessary. Using the shooting method described above, we predict the spatial profiles at steady state for bile pressure, velocity and solute concentration as shown in Fig. 5 and Extended Data Fig. 9. The obtained solutions were qualitatively in agreement with the asymptotic state in numerical simulations of the time-dependent problem along a single tube (data not shown) which we studied using the software Morpheus[17].

### Bile acids and serum markers

Individual serum bile acids and their precursors were analysed either by gas-liquid chromatography mass spectrometry (GLC-MS) or liquid chromatography (HPLC) as described[18].

### Statistical analysis

For the spatially-resolved quantifications (i.e. statistics along the CV-PV axis), the mean ± SEM values per zone are shown. Global quantifications (i.e liver enzymes, bile acids and overall tissue parameters) are shown using box-plots. Whereas median values were shown as red lines, the 25th and 75th percentiles were represented by the blue bottom and top edges of the boxes, respectively. The whiskers were extended up to the most extreme data points that are not considered outliers. The statistical significance analysis was performed using either two-sided (for liver enzymes and bile acids) or one-sided (tissue parameters) t-tests assuming unequal variances. All statistical analysis were performed using MATLAB2018b.
